# *N*-glycosylation modulates enzymatic activity of *Trypanosoma congolense* trans-sialidase

**DOI:** 10.1101/2021.12.13.472379

**Authors:** Jana Rosenau, Isabell Louise Grothaus, Yikun Yang, Nilima Dinesh Kumar, Lucio Colombi Ciacchi, Sørge Kelm, Mario Waespy

**Affiliations:** University of Bremen, Centre for Biomolecular Interactions Bremen, Faculty for Biology and Chemistry, 28359 Bremen, Germany; University of Bremen, Hybrid Materials Interfaces Group, Faculty of Production Engineering, Bremen Center for Computational Materials Science, Center for Environmental Research and Sustainable Technology (UFT), and MAPEX Center for Materials and Processes, 28359 Bremen, Germany; Innovation Research Institute of Traditional Chinese Medicine, Shandong University of Traditional Chinese Medicine, Jinan 250355, Shandong, China; Department of Biomedical Sciences of Cells and Systems, University Medical Center Groningen, University of Groningen, 9712 CP Groningen, Netherlands

**Keywords:** proteoglycan, molecular dynamics, mass spectrometry (MS), enzyme kinetics, circular dichroism, *N*-glycosylation, protein-glycan interactions, trans-sialidases, *Trypanosoma*

## Abstract

Trypanosomes cause the devastating disease trypanosomiasis, in which the action of trans-sialidase (TS) enzymes harbored on their surface is a key virulence factor. TS are *N*-glycosylated, but the biological functions of their glycans has remained elusive. In this study, we investigated the influence of *N*-glycans on the enzymatic activity and structural stability of TconTS1, a recombinant TS from the African parasite *Trypanosoma congolense*. The enzyme was expressed in CHO Lec1 cells, which produce high-mannose type *N*-glycans similar to the TS *N*-glycosylation pattern *in vivo*. MALDI-TOF MS data revealed that up to eight putative *N*-glycosylation sites were glycosylated. *N*-glycan removal via EndoH_f_ treatment of TconTS1 led to a decrease in substrate affinity relative to the untreated enzyme, but apparently has no impact on the conversion rate. No changes in secondary structure elements of hypoglycosylated TconTS1 were observed in circular dichroism experiments. Molecular dynamics simulations provided evidence for interactions between monosaccharide units of the highly flexible *N*-glycans and some conserved amino acids located at the catalytic site. These interactions led to conformational changes, possibly enhancing substrate accessibility and enzyme-substrate complex stability. The here-observed modulation of catalytic activity via *N*-glycans represents a so far unknown structure-function relationship potentially inherent in several members of the TS enzyme family.

## Introduction

Glycosylation, or the covalent attachment of carbohydrates to the polypeptide chain of proteins, is the most diverse protein modification in eukaryotic cells. One type of protein glycosylation is the *N*-linked glycosylation, the attachment of an oligosaccharide to an asparagine residue (*N*-glycan) in a specific protein sequence. *N*-glycans are branched, tree-like polysaccharide structures built up of different monosaccharide units, which are linked via different glycosidic bonds.

*N*-glycans are involved in folding and stability of proteins, regulate protein functions, represent target structures for lectins or antibodies, and can function as mediators of cell-matrix interactions and cell-cell recognition and communication (1, 2). Recent studies of *N*-glycans include molecular dynamics simulations of the SARS-CoV-2 spike protein’s glycan shield, the characterization of the glycosylated infectious bronchitis virus spike protein as well as the heavily glycosylated receptor CD22 on B-cells by means of wet lab-based approaches (3–5).

In this study, we investigated the influence of *N*-glycans on the catalytic transfer activity, thermal stability and structural conformation of *Trypanosoma congolense* trans-sialidase (TconTS) 1, by determining enzyme kinetics, mass spectrometry (MS) measurements and molecular dynamics simulations. TS are unusual enzymes expressed by different species of the parasitic genus *Trypanosoma* (*T*.). This includes human pathogens causing Chagas disease in South America (*T. cruzi*) and sleeping sickness (Human African Trypanosomiasis, HAT) in Africa (*T. brucei*), as well as animal pathogens being responsible for the Animal African trypanosomiasis (AAT) (mainly *T. brucei brucei, T. congolense, T. vivax*) (6–9). The glycosylphosphatidylinositol (GPI)-anchored enzymes preferentially transfer α2,3-linked sialic acids (Sia) from host-cell glycoconjugates to terminal β-galactose residues of glycoproteins present on the parasite’s surface, thus creating a new α2,3-glycosidic linkage (6, 7, 10). This surface sialylation has different beneficial functions for the parasite, which is unable to synthesize Sia *de novo*. In particular it promotes survival in the insect vector and enables to escape the host’s immune system (11–15). Thus, TS are important virulence factors and represent promising drug targets and vaccine candidates to combat the fatal diseases caused by trypanosomes.

The enzymatic mechanisms and biological functions of TS have been under study for many years, although little attention has been drawn to the influence of post-translational modifications such as glycosylation (16–21).

Interestingly, the presence of high-mannose type *N*-glycans has been indicated for many TS indirectly by concanavalin A (ConA) purification (7, 22, 23). On the other hand, hybrid or complex *N*-glycans have not been reported for TS. It has been postulated that *N*-glycans are involved in TS oligomerization (24). However, its influence on enzyme activity is still debated (16, 17, 20, 21). Previous experiments were performed with recombinant TS expressed by *Pichia (P.) pastoris*, which produces hypermannosylated *N*-glycans (20, 21). However, trypanosomal surface proteins were reported to harbor shorter high-mannose type *N*-glycans (25–30). No differences in Sia transfer activity were observed between glycosylated and deglycosylated recombinant TS expressed by *P. pastoris* (20, 21) even though kinetic parameters were not determined.

Recombinant TconTS1 from the animal pathogen *T. congolense* expressed in CHO Lec1 cells was used in this study (31). Natively, TconTS1 is expressed by procyclic insect-infective trypanosomes as well as bloodstream-form trypanosomes in mammalian hosts, and is involved in desialylation of erythrocytes, which contributes to anemia (32, 33). Like all TS, TconTS1 consists of an *N*-terminal catalytic domain responsible for the transfer of Sia, and of a C-terminal lectin-like domain whose biological function remains unclear. Recently, we demonstrated that the TconTS lectin-like domain can modulate enzymatic activity (34). The catalytic and lectin-like domain are connected via an α-helix (Fig. 1) (16, 17, 31). The *N*-terminus includes a signal sequence, whereas the C-terminus comprises a potential GPI anchor attachment site (31). So far, 17 TconTS-like genes have been described for *T. congolense,* from which 11 can be grouped into the TconTS1 family due to their high amino acid sequence identity (>96 %). TconTS1b was chosen in this study, because it possesses one of the highest enzyme activity among TconTS family (31, 35) and is simply called TconTS1 onwards.

**Figure 1.**
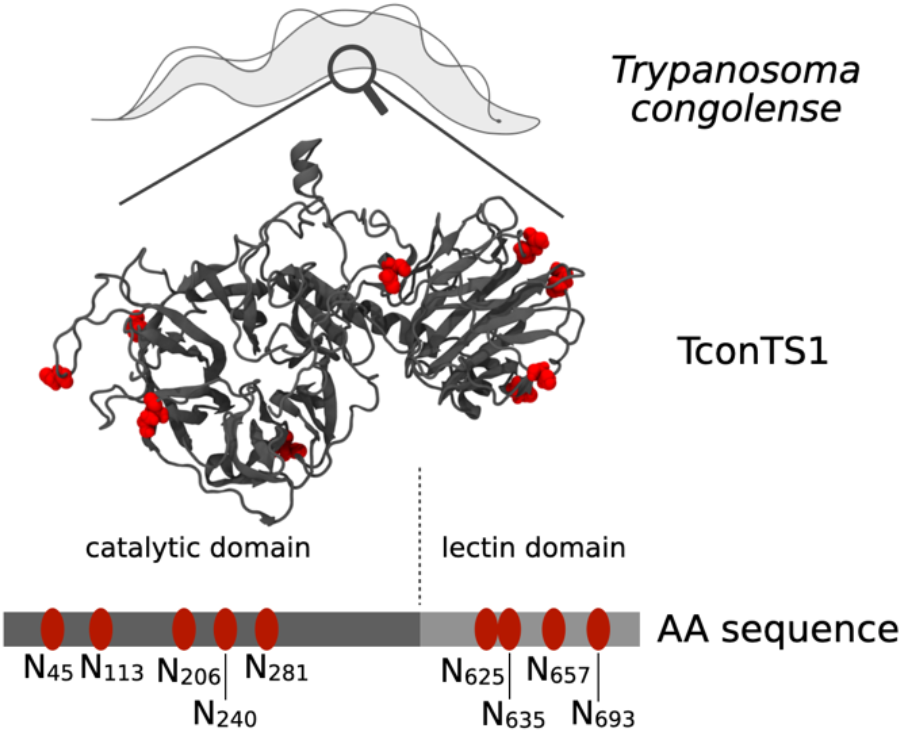
Molecular model of TS originating from *Trypanosoma congolense* (TconTS1) with *N*-glycosylation sites. Asparagine residues in the motif N-X-S/T as putative *N*-glycosylation sites are highlighted in red. The positions of each asparagine are labelled in the amino acid sequence and sorted into the catalytic domain (CD) or the lectin-like domain (LD).

An important structural feature of TconTS1 are the predicted nine *N*-glycosylation recognition sequences (Fig. 1), distributed across both the catalytic and lectin-like domains (31). It is noteworthy that TS sequences from African trypanosomal species contain a higher number of putative *N*-glycosylation sites (6–9) compared to those of species from South-American trypanosomes (2–5) including the structural closely related sialidase (SA) from *T. rangeli* (9, 20, 31, 35–38). Crystallization of the SA from *T. rangeli* revealed that all five potential *N*-glycosylation sites were occupied with *N-*glycans, although only the innermost monosaccharide could be detected, giving no hint about the type of *N*-glycosylation (16). The only crystallized TS from *T. cruzi*, however, was a recombinant protein containing several mutations in the amino acid sequence (17). Thus, the *N*-glycosylation pattern of this TS remains unresolved. In contrast to *T. brucei* (39), there are no database entries on potential oligosaccharyltransferase isoforms in *T. congolense*. Therefore, no further assessment of potential glycosylation pattern and glycan donor- and peptide-acceptor specificity can be made.

To investigate *N*-glycosylation-dependent structure-function effects, we expressed recombinant TconTS1 in leuco-phytohemagglutinin (L-PHA)-resistant Lec1 Chinese hamster ovary (CHO) cells (40, 41). This *N*-glycosylation mutant cell line is i) unable to synthesize complex and hybrid *N*-glycans and ii) consequently accumulates high-mannose type *N*-glycans of the composition Man_5-9_GlcNAc_2_, which mimics the situation reported for African trypanosomes preferentially synthesizing oligosaccharides bearing Man_5-9_GlcNAc_2_ *N*-glycans (25–30, 40, 41). For these reasons, we have selected CHO Lec1 cell line as a *N*-glycosylation model system in this study. The presence and composition of *N*-glycans at the putative *N*-glycosylation sites in TconTS1 was analyzed qualitatively by matrix assistant laser desorption ionization – time of flight (MALDI-TOF) mass spectrometry (MS), in order to identify modified sites and oligosaccharide distribution. Enzyme activities of glycosylated and Endoglycosidase H (EndoH_f_) treated TconTS1 were analyzed via quantification of reaction products using high performance anion exchange chromatography (HPAEC) with pulsed amperometric detection (PAD). Circular Dichroism spectroscopy was used to compare the secondary structures and stabilities of glycosylated versus EndoH_f_-treated TconTS1. Finally, molecular dynamics (MD) simulations were performed to characterize potential protein-glycan and glycan-glycan interactions altering the dynamics of amino acids in close proximity to the catalytic domain.

## Results

### 1. Mapping the *N*-glycosylation pattern of TconTS1

To characterize the *N*-glycosylation pattern of TconTS1 in detail, we expressed TconTS1 in CHO Lec1 cells (Fig. S1A, S2) (31). High-mannose type *N*-glycans of the recombinant protein were detected by ConA lectin blots (Fig. 2A, S3A/B). For the selective removal of *N*-glycans, TconTS1 was treated with EndoH_f_. A clear band shift of approximately 10 kDa for TconTS1 after 4 hours of incubation with EndoH_f_ was observed, indicating a size reduction as a consequence of the removal of *N*-glycans from TconTS1 (Fig. 2A, S4A/B). In addition, less binding of ConA to EndoH_f_ treated TconTS1 was observed by lectin blots, indicating a loss of *N*-glycan structures (Fig. 2A lower panel). However, deglycosylation was not complete after 4 hours of incubation and even further addition of EndoH_f_ or increased incubation times to up to 48 hours did not result in complete deglycosylation. EndoH_f_-treated TconTS1 incubated overnight (16 hours) (Fig. S4C/D) was used for *N*-glycosylation pattern analysis in this study. This modified TconTS1 is termed hypoglycosylated TconTS1 (H-TconTS1) onwards. Raw data and further details of western and lectin blot analysis are given in Section S2 of the Supporting Information (SI).

**Figure 2.**
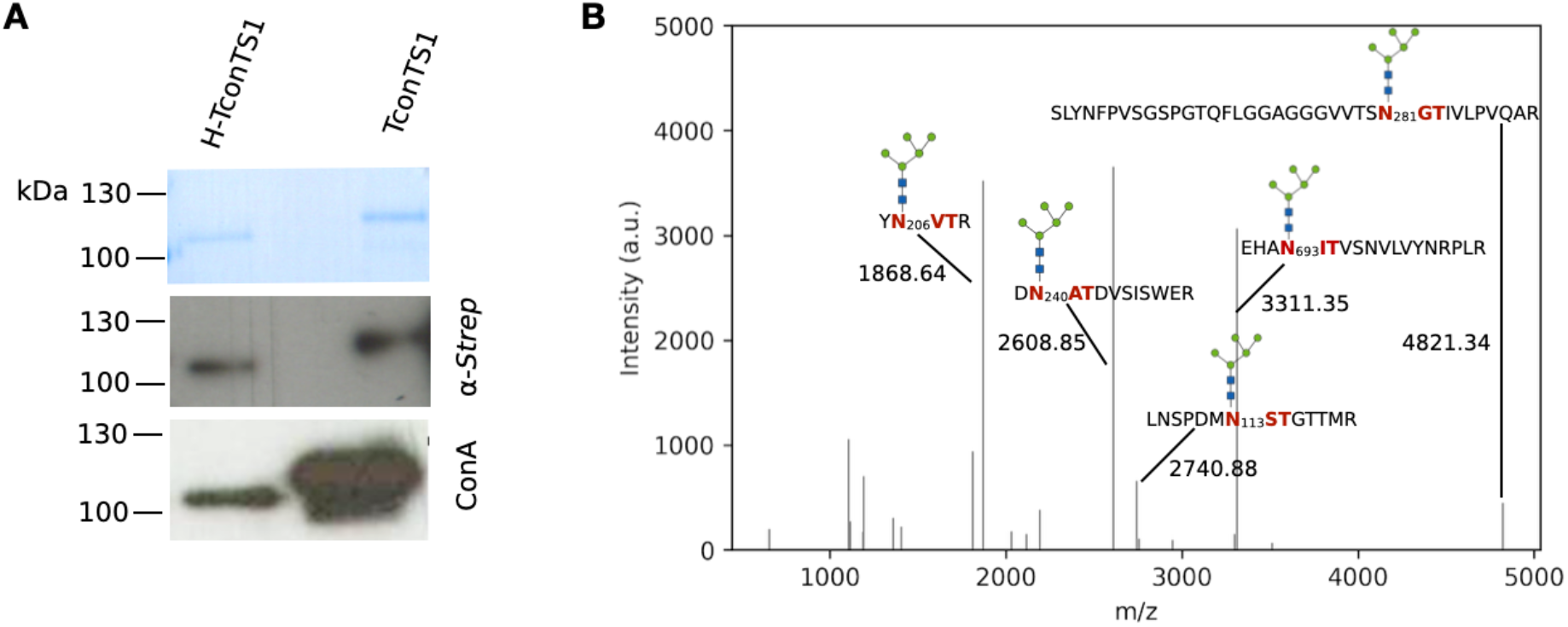
Mapping the *N*-glycosylation profile of untreated (TconTS1) and hypoglycosylated TconTS1 (H-TconTS1). A) TconTS1 and H-TconTS1 (deglycosylated for 4 h) were analyzed by SDS-PAGE with subsequent Coomassie staining (upper panel with 600 ng of protein), western blot analysis using an anti-*Strep*-tag antibody (middle panel with 400 ng of protein) and lectin blotting using ConA (lower panel with 100 ng of protein). Exposure time of western blot is 5 sec and 60 sec for the ConA blot. Results indicate the presence of high-mannose type *N*-glycans. B) MALDI-TOF MS analyses were performed to identify *N*-glycosylation sites of TconTS1. Glycopeptides from protease-digested TconTS1 were ConA-purified to concentrate glycopeptides and reduce the spectrum complexity. Peak lists were extracted from MALDI-TOF mass spectra, plotted with python and annotated with corresponding masses and glycopeptide fragments, respectively. Monosaccharide symbols follow the Symbol Nomenclature for Glycans (SNFG) (83).

MALDI-TOF MS experiments were performed to identify *N*-glycan structures and distribution pattern. In order to analyze TconTS1 and H-TconTS1 by MALDI-TOF MS, both enzymes were proteolytically digested to shorter peptide and glycopeptide fragments. Whenever an asparagine residue in the N-X-S/T motif of a certain glycopeptide is glycosylated, the m/z ratio increases by exactly the mass of the conjugated *N*-glycan relative to the non-glycosylated peptide.

The setup of the Bruker Autoflex Speed device used in this study does not offer in-source fragmentation. Thus, we employed different analyses with varying sample preparations to cross-validate identified *N*-glycans of TconTS1. First, we used two different proteases, trypsin and chymotrypsin, to generate glycopeptides. An advantage of this strategy is that treatment with either trypsin or chymotrypsin results in different glycopeptide profiles due to their different specific protease recognition sites (for trypsin digestion see Fig. S5B). As a result, the same potential *N*-glycosylation site can be found within different glycopeptides after either trypsin or chymotrypsin digestion. Second, glycopeptides were purified using ConA sepharose to specifically enrich and concentrate glycopeptides comprising high-mannose type *N-*glycans, and as a consequence to lower the complexity of spectra (Fig. 2B, S5A). Third, H-TconTS1 was analyzed using the same strategy described above. EndoH_f_ cleaves within the chitobiose core of high-mannose type *N-*glycans, leaving a residual GlcNAc at the glycopeptide. Thus, peptides generated from H-TconTS1 that comprise a putative *N-*glycosylation site and exhibit a mass difference of m/z 203.08 (one HexNAc, corresponding to one GlcNAc) relative to the non-glycosylated peptide indicate the presence of a high-mannose type *N*-glycan at that site (Fig. S5C). A summarizing evaluation of these different approaches is given in Table 1.

**Table 1:** MALDI-TOF MS analyses of the N-glycan profile of TconTS1. For analysis, the untreated TconTS1 (glycosylated, G) and the EndoHf treated H-TconTS1 (H) were digested with trypsin or chymotrypsin. Spectra were analyzed for masses corresponding to glycopeptides with high-mannose type N-glycans and for non-glycosylated peptides with potential N-glycosylation sites. Spectra of H-TconTS1 were additionally analyzed for glycopeptides with HexNAc residues since a residual GlcNAc remains attached to the protein N-glycosylation sites after EndoHf treatment. In another approach, glycopeptides from both proteins were purified with ConA after protease digestion and spectra were analyzed for masses of peptides with high-mannose type N-glycans. *Glycopeptide mass was only detected once during multiple analyses.

|  | ***N*-glycan** | **N<sub>45</sub>** | **N<sub>113</sub>** | **N<sub>206</sub>** | **N<sub>240</sub>** | **N<sub>281</sub>** | **N<sub>625</sub>** | **N<sub>635</sub>** | **N<sub>657</sub>** | **N<sub>693</sub>** |
| --- | --- | --- | --- | --- | --- | --- | --- | --- | --- | --- |
| Trypsin | Non-glycosylated |  |  |  | **G** |  |  |  |  |  |
|  | Man<sub>5</sub> GlcNAc<sub>2</sub> |  | **G** | **G** | **G** | **G** |  |  |  | **G** |
|  | Man<sub>6</sub> GlcNAc<sub>2</sub> |  |  |  |  |  |  |  |  |  |
|  | Man<sub>7</sub> GlcNAc<sub>2</sub> |  |  |  |  |  |  |  |  |  |
|  | Man<sub>8</sub> GlcNAc<sub>2</sub> |  |  |  |  |  |  |  |  |  |
|  | Man<sub>5</sub> GlcNAc<sub>2</sub> Fuc<sub>1</sub> |  |  | **G** |  |  |  |  |  |  |
|  | HexNAc |  | **H** | **H** | **H** | **H\*** |  |  |  | **H** |
| Trypsin + ConA | Man<sub>5</sub> GlcNAc<sub>2</sub> |  | **G** | **G/H** | **G** | **G** |  |  |  | **G** |
|  | Man<sub>6</sub> GlcNAc<sub>2</sub> |  |  | **G/H** |  |  |  |  |  | **H\*** |
|  | Man<sub>7</sub> GlcNAc<sub>2</sub> |  |  | **G** |  |  |  |  |  |  |
|  | Man<sub>8</sub> GlcNAc<sub>2</sub> |  |  | **H** |  |  |  |  |  |  |
|  | Man<sub>5</sub> GlcNAc<sub>2</sub> Fuc<sub>1</sub> |  |  |  |  |  |  |  |  |  |
| Chymo-<br>trypsin | Non-glycosylated |  | **G** | **G** |  |  | **G/H** |  |  | **H** |
|  | Man<sub>5</sub> GlcNAc<sub>2</sub> | **G** | **H\*** | **G** |  |  |  |  |  |  |
|  | Man<sub>6</sub> GlcNAc<sub>2</sub> |  | **G\*** |  |  |  |  |  |  |  |
|  | Man<sub>7</sub> GlcNAc<sub>2</sub> |  |  |  |  |  |  |  |  |  |
|  | Man<sub>8</sub> GlcNAc<sub>2</sub> |  |  |  |  |  |  |  |  |  |
|  | Man<sub>5</sub> GlcNAc<sub>2</sub> Fuc<sub>1</sub> |  |  |  |  |  |  |  |  |  |
|  | HexNAc | **H** |  | **H** |  |  |  |  | **H** | **H** |
| Chymo-<br>trypsin +<br>ConA | Man<sub>5</sub> GlcNAc<sub>2</sub> | **G** |  | **G/H** |  |  |  |  | **G** |  |
|  | Man<sub>6</sub> GlcNAc<sub>2</sub> |  |  | **G/H** |  |  |  |  |  |  |
|  | Man<sub>7</sub> GlcNAc<sub>2</sub> |  |  |  |  |  |  |  |  |  |
|  | Man<sub>8</sub>GlcNAc<sub>2</sub> |  |  | **G/H** |  |  |  |  |  |  |
|  | Man<sub>5</sub>GlcNAc<sub>2</sub>Fuc<sub>1</sub> |  |  |  |  |  |  |  |  |  |

Glycopeptides comprising high mannose-type *N*-glycans were detected especially in the catalytic domain. *N*-glycans attached to N113 and N206 were detected in both trypsin-treated and chymotrypsin-treated TconTS1, (Table 1). The mass difference corresponding to an *N-*glycan of the composition Man_5_GlcNAc_2_ was predominantly detected at N113, whereas N206 also exhibited the composition Man_6-8_GlcNAc_2_ (Table 1 and Table S2). In addition, when analyzing H-TconTS1, HexNAc residues were detected at positions N113 and N206. But for both sites also corresponding non-glycosylated peptides were detected. Interestingly, *N*-glycosylation at N206 was still detectable after 16 h of EndoH_f_ treatment. This finding is supported by our ConA lectin blot results, demonstrating the binding of ConA to H-TconTS1 at long exposure times (Fig. 2 and Fig. S4).

Glycopeptides containing high-mannose type *N*-glycans conjugated to other *N*-glycosylation sites were either detected in trypsin- or chymotrypsin-treated samples. For example, the mass difference corresponding to the composition Man_5_GlcNAc_2_ was identified at N45 and N657 (chymotrypsin) and at N240, N281 and N693 (trypsin) (Table 1). In addition, the same glycopeptides were identified in the corresponding ConA-purified samples (Table 1). When analyzing H-TconTS1, glycopeptides with mass differences corresponding to a residual HexNAc residue were detected in the glycopeptides obtained after digestion with the corresponding proteases (Table 1). N240 was also detected in its non-glycosylated form as well as N693 (only in H-TconTS1), indicating a heterogeneity at this site. Peptides containing N625 were only found as the non-glycosylated form. Additional details regarding the performed MALDI-TOF MS analysis can be found in Section S3 of the SI.

In summary, the compiled data has revealed *N*-glycosylation predominantly in the catalytic domain of recombinant TconTS1 allowing qualitative prediction of the predominant *N*-glycan composition for different *N*-glycosylation sites.

### 2. *N*-glycosylation of TconTS1 influences the affinity to the Sia acceptor substrate in the transfer reaction

To test the impact of *N*-glycosylation on enzyme activity, we performed activity assays using H-TconTS1 and TconTS1. For these assays TconTS1 was exposed to identical conditions applied for the EndoH_f_ treatment. Fetuin and lactose were used as Sia donor and acceptor substrates in enzyme reactions, as previously described (31, 35). The transfer reaction product 3’sialyllactose (3’SL) was quantified by HPAEC-PAD. This HPLC-based method allows for the separation and detection of the educt lactose, and the product 3’SL. Incubation times of 30 minutes and 600 µM fetuin-bound Sia were used in these experiments. A 600 µM concentration of fetuin-bound Sia was used previously to determine kinetic parameters for TconTS, allowing us to compare our results with that available from the literature (31). The Sia concentration used corresponds approximately to one third of the Sia concentration reported to be present on glycoproteins in blood serum (42, 43). In human blood serum a high proportion of protein-bound Sia is α2,6-linked (44, 45). A lactose concentration series was applied in order to calculate the corresponding Michaelis-Menten kinetic parameters K_M_ and V_max_.

Two different TconTS1 preparations (biological duplicates) and the corresponding hypoglycosylated proteins were investigated. Both data sets revealed that H-TconTS1 produced lower amounts of 3’SL relative to TconTS1 in these experiments (Fig. 3A). Interestingly, the calculated V_max_ for the Sia acceptor substrate lactose of about 2.4 µmol 3’SL/(min x mg enzyme) for data set 1 and to 4.1 µmol 3’SL/(min x mg enzyme) for data set 2 was found to be identical within the error range for both data sets (Fig. 3B). These results indicate that the 3’SL production rate of TconTS1 at saturated Sia acceptor concentrations is not influenced by its *N*-glycosylation state. Differences in V_max_ values between the data sets might be explained by varying amounts of active enzyme after purification. However, to study the influence of *N*-glycosylation on enzyme activity, the kinetic parameters were calculated separately for each data set.

**Figure 3.**
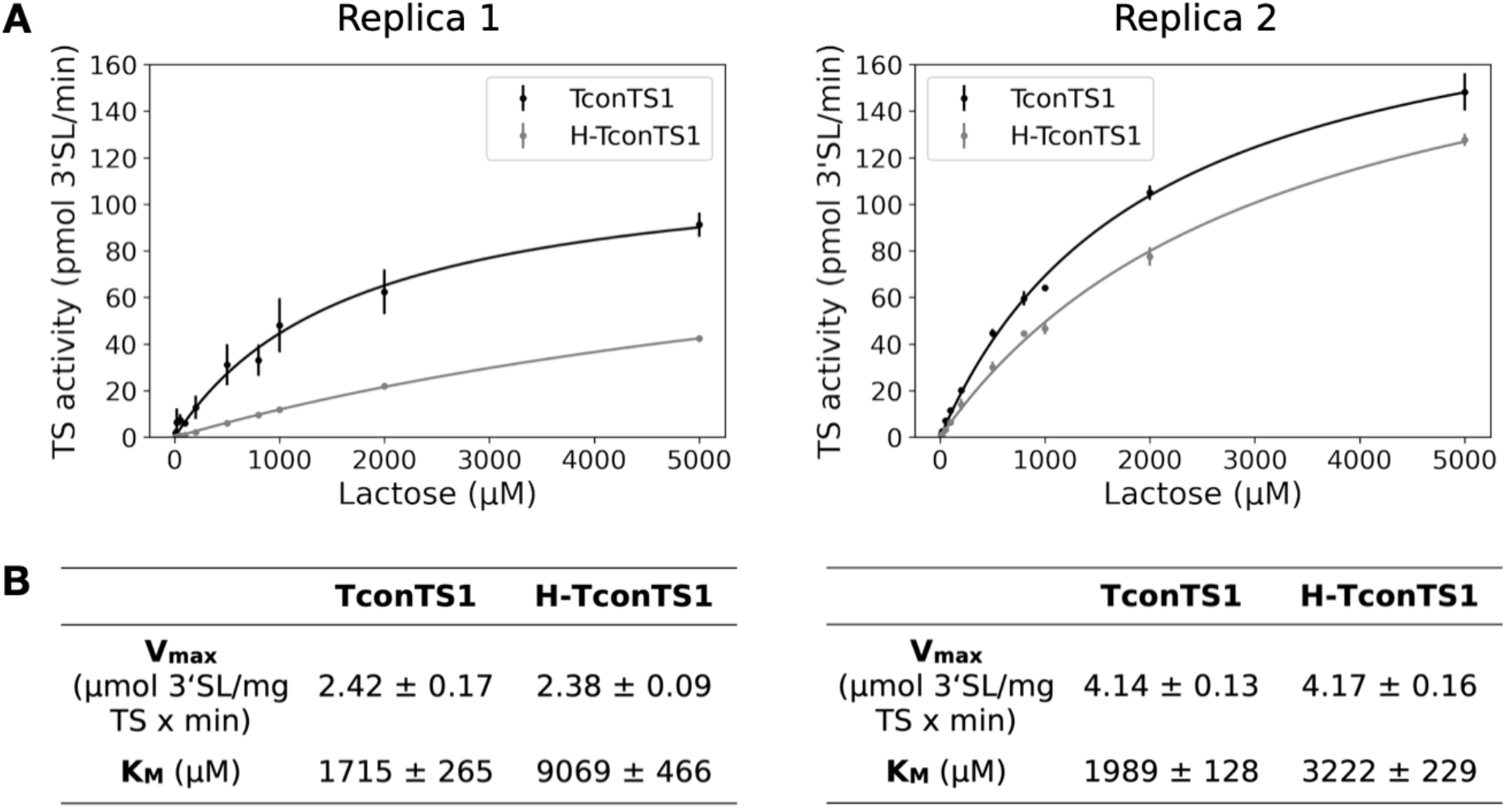
EndoHf treated H-TconTS1 shows lower K_M_ compared to TconTS1. A/B) TS activities for TconTS1 and H-TconTS1 were determined using fetuin as Sia donor and a lactose concentration series as Sia acceptor. Production of 3’sialyllactose was monitored and Michaelis-Menten kinetic parameters apparent KM and Vmax for lactose were evaluated using SigmaPlot11. Data points are means ± standard deviation of three technical replicas for each biological replica.

Interestingly, the K_M_ values for lactose differed between TconTS1 (1.7 and 2.0 mM) and H-TconTS1 (9.0 and 3.2 mM), indicating a 1.6- to 5-fold lower Sia acceptor substrate affinity for H-TconTS1 relative to TconTS1 (Fig. 3). The K_M_ values determined in this study are similar to that published by Koliwer-Brandl et al. (31) (1.7 mM). Small variations in the K_M_ of H-TconTS1 between both data sets might be a result of differences in the number of *N*-glycans remaining on these hypoglycosylated TconTS1 preparations. Since MALDI-TOF MS does not reveal single protein-resolved *N*-glycosylation pattern, a comparison of spectra generated from H-TconTS1 peptides of both data sets 1 and 2 has not be performed.

In summary, the same trend was observed for both data sets: a significant decrease in the K_M_ value for lactose after EndoHf treatment, while the V_max_ value remained constant.

### 3. EndoHf treatment of TconTS1 does not alter the overall secondary structure

Once we were able to determine TconTS1’s *N*-glycan profile and its influence on the enzyme activity, we addressed the question, whether removal of *N*-glycans affects enzyme stability. Since the presence of *N*-glycans can impact the protein’s function via altering the protein structure (46), we performed circular dichroism experiments to investigate the influence of *N*-glycans on TconTS1’s secondary structure stability.

Circular dichroism spectra of TconTS1 and H-TconTS1 were analyzed under similar conditions used for the enzyme activity measurements (35°C) (Fig. 4A). The recorded spectra do not show any significant difference over the recorded wavelength range, indicating that both enzymes share the same common secondary structure. In fact, calculated secondary structural elements were identical in both cases with 37% of β-sheets, 13% of α-helices, 11% of turns and 39% of unstructured (other) components in both cases (Table S4).

**Figure 4.**
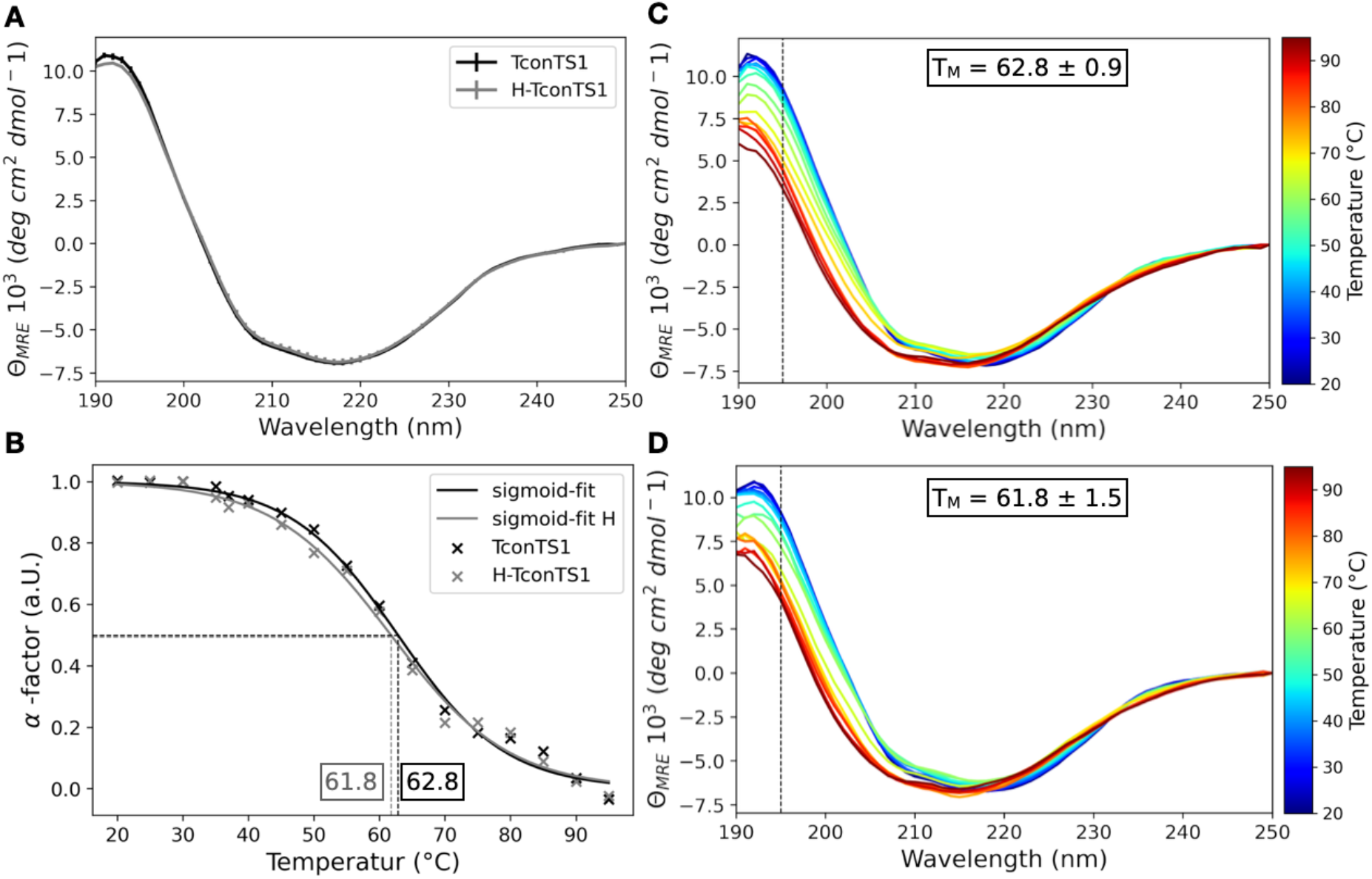
Influence of *N*-glycans on TconTS1 secondary structure and stability. A) Circular dichroism spectra of untreated TconTS1 and EndoHf treated H-TconTS1 were measured at 35°C in 10 mM phosphate buffer pH 7.4 (means ± standard deviation of two biological replicas). B) The midpoint of unfolding (TM = α-factor = 0.5) from a folded state (α-factor ∼ 1.0 at 20 °C) to a partially unfolded intermediate (α-factor ∼ 0.0 at 95 °C) were determined for TconTS1 and H-TconTS1 by fitting a sigmoid function to the data. Circular dichroism spectra of temperature-ramping experiments with TconTS1 (C) and H-TconTS1 (D) were recorded in 5°C steps and a 5 min equilibration time at each step. The midpoint of unfolding (TM) is determined at 195 nm (dashed line) at the flex point of a sigmoidal function fitting the temperature curve.

Spectra recorded during temperature-ramping experiments revealed a high heat stability for both protein preparations, as TconTS1 and H-TconTS1 still kept intact secondary structural elements up to 95°C (Fig. 4C/D). The variation of the spectra intensity in the range between 190 and 210 nm during heating indicates a partial unfolding, taking place between 60°C and 70°C, with a melting temperature (T_M_) of about 62°C for both proteins (Fig. 4B). Thus, TconTS1 and H-TconTS1 exhibit the same secondary element distribution and the same heat stability (as quantified by T_M_ for partial unfolding).

In summary, the circular dichroism experiments did not provide evidence for an altered overall secondary structure of H-TconTS1 as explanation for its lower substrate affinity observed. However, *N*-glycosylation-induced changes of the tertiary structure remain as possible explanation because they are not detectable in circular dichroism spectra in the wavelength region measured.

### 4. Structural importance of TconTS1 *N*-glycans indicated by molecular dynamics simulations

#### 4.1 The *N*-glycan shield

Molecular dynamics simulations can facilitate the study of mechanisms at the atomistic level and were performed to obtain an in-depth picture of *N*-glycan-protein interactions in TconTS1. Due to the lack of experimentally derived structures for TconTS enzymes, homology models were constructed on the basis of *T. rangeli* sialidase (TranSA) and *T. cruzi* TS (TcruTS). Further details about the model construction are given under Experimental procedures (see also Section S5 of the Supporting information). Although TconTS1 only shares an amino acid sequence identity of 48 % with TcruTS, both enzymes reveal an high overall structural similarity (Fig. S6) and only differ in three amino acids reported to be important for TS activity (31). In order to generate a *N*-glycosylated structural model of TconTS1, Man_5_GlcNAc_2_ glycans were attached at positions N45, N113, N206, N240, N281 and N693, as identified by our MALDI-TOF MS experiments. In this model, all potential *N*-glycosylation sites are glycosylated in the catalytic domain, whereas only one out of four potential sites is glycosylated in the lectin-like domain (Fig. S7A). In the model used to simulate H-TconTS1, single GlcNAc residues were attached to positions that were glycosylated in TconTS1, mimicking the residual monosaccharide after EndoHf treatment (Fig. S7B). On purpose, a TconTS1 model harboring all *N*-glycosylation sites identified in this study (Table 1) and a corresponding fully deglycosylated model with remaining GlcNAc (H-TconTS1) were employed.

At first, standard MD simulations were performed for TconTS1 (Fig. S7A) and H-TconTS1 (Fig. S7B) without bound substrates, in order to observe the dynamical behavior of the covalently linked *N*-glycans. Six out of nine asparagine residues in the motif N-X-T/S are located at the tail of loop regions (Fig. 5A), which are mostly part of turns or coils and framed by β-sheet regions. Results of our in-depth analysis can be found in the SI section S5 in Figure S8. The terminal position and flexibility of these structural elements allow for large motion amplitudes and internal flexibility of the *N*-glycan trees. These movements enable interactions among glycans in structural proximity, for instance intermolecular hydrogen bonds between the *N*-glycans at positions N113 and N240 (Fig. 5B), notwithstanding their distance in the protein sequence. Furthermore, an overlay of the averaged glycan distribution recorded every 5 ns during the simulation (Fig. 5C/D) revealed a dense glycan coverage (shielding) of the protein, especially for the catalytic domain, but except the direct entrance to the active site (Fig. 5C).

**Figure 5.**
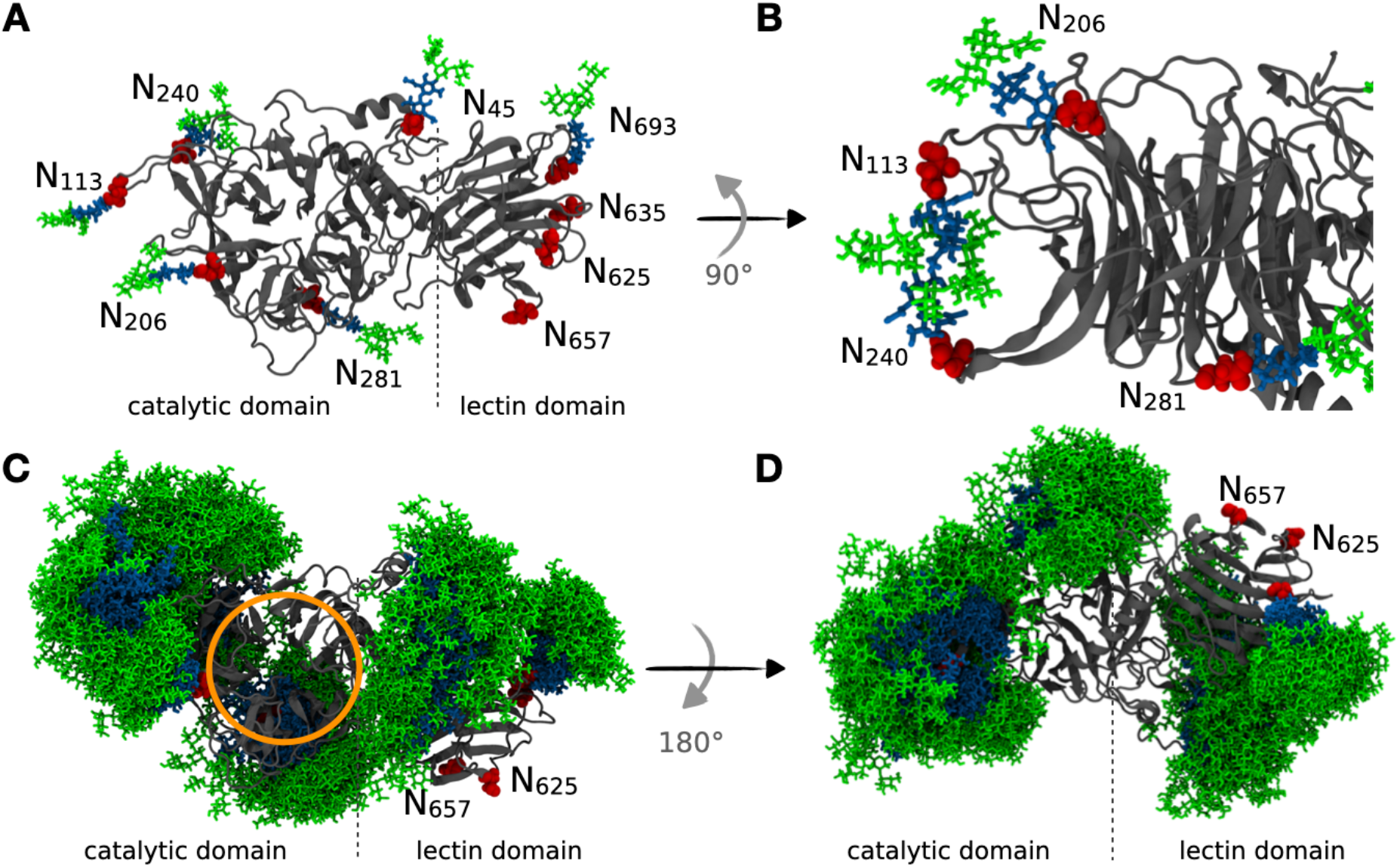
Analysis of the dynamics of TconTS1’s *N*-glycan shield. A) Atomistic model of TconTS1 with attached Man5GlcNAc2 *N*-glycans (Man: green, GlcNAc: blue) at the asparagine residues (red) identified in MALDI-TOF MS experiments. B) Interactions between N240 and N113 glycans mediated by hydrogen bonds observed during the MD simulations. C) Overlay of all *N*-glycan positions recorded every 5 ns over a simulation time of 500 ns, with the protein backbone (grey) aligned in all frames and the active site indicated by an orange circle. D) Same as C, with the protein rotated by 180°. The C-terminal SNAP-*Strep* is not shown in all structures.

#### 4.2 Dynamics of conserved amino acids in the catalytic domain of substrate-free TconTS1

Interactions of *N*-glycans with highly conserved amino acids being essential for the catalytic activity of TconTS1 (31, 47) were analyzed over the 500 ns MD trajectories. We especially focused on D150, E324 and Y438 residues known to be directly involved in catalysis as well as R126, R144, Y211, W212, R339, Y408 and R410, which are involved in substrate binding.

In TconTS1, D150 was observed to shift from the interior of the active site (Fig. 6A) towards an exterior position (Fig. 6B), increasing its distance from R410, which was stationary within the active site (Fig. 6C). Interestingly, this shift seems to be stabilized by a hydrogen bond formed between D150 and the *N*-glycan attached to N206 (Fig. 6C/D). Further detailed analysis revealed that this process was initiated by hydrogen bond formation between Y151 and the *N*-glycan at N206, already leading to a partial shift of D150 and making it more accessible to interact with the *N*-glycan (Fig. 6C). In contrast, for H-TconTS1 D150 was found to be by far less mobile (Fig. 6C).

**Figure 6.**
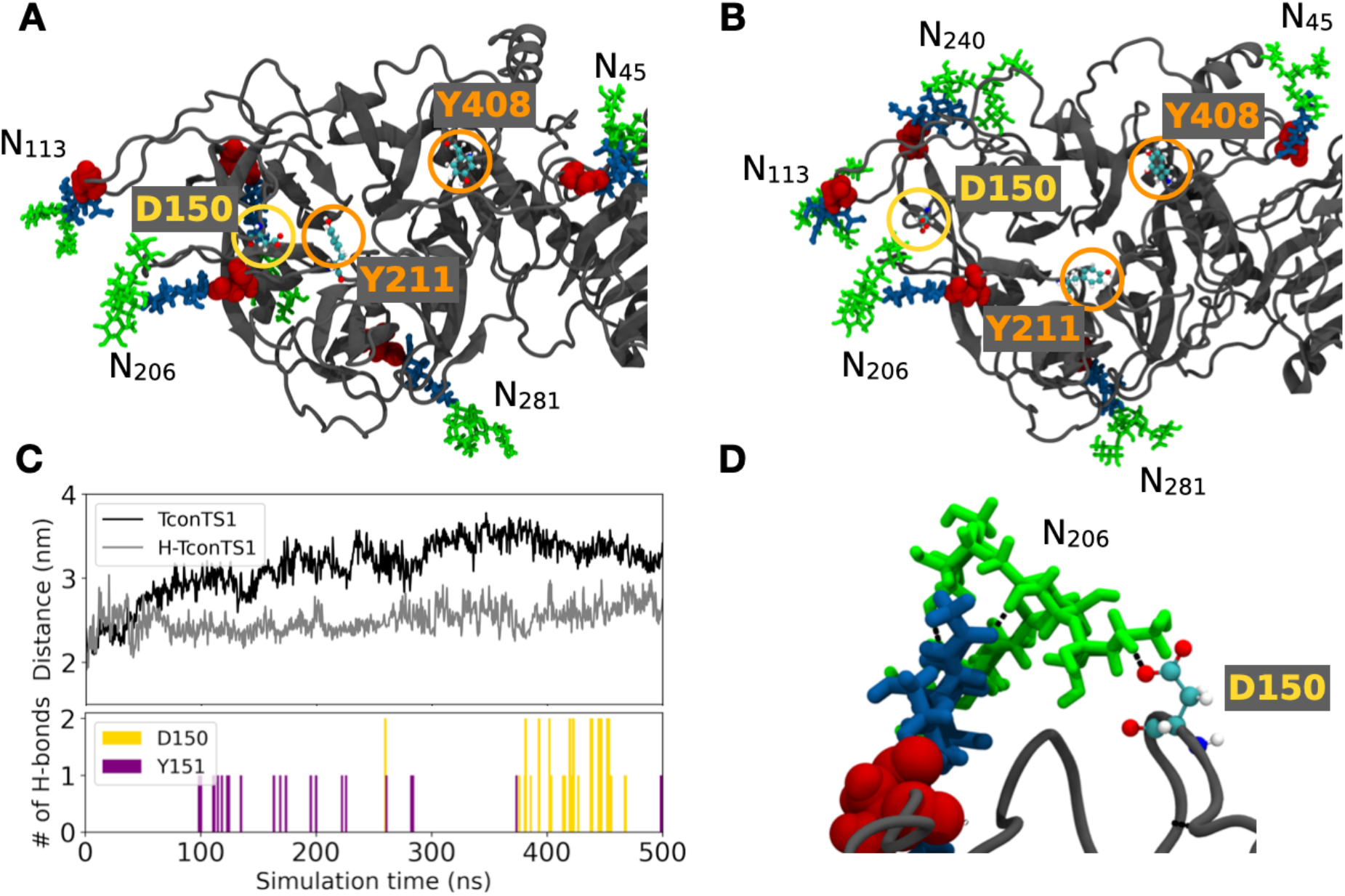
Protein-glycan interactions drive active site rearrangements observed in MD simulations of TconTS1 without substrate. A) Amino acids of the catalytic site were in close proximity at the beginning of the simulation (Snapshot at 100 ns). B) D150 moved out of the catalytic site and formed hydrogen bonds with glycan N206 until the end of the simulation (Snapshot at 350 ns). C) Time evolution of the distance between the center of D150 and the center of R410 for TconTS1 (black) and H-TconTS1 (grey) as well as numbers of hydrogen bonds for TconTS1 between glycan N206 and D150/Y151. For H-TconTS1, no hydrogen bonds were observed. D) Detail of the hydrogen bonds (black dashed lines) between D150 and glycan N206 at its terminal mannose branches. D150 is circled in yellow and the ligand-binding residues Y211 and Y408 are circled in orange and represented in ball-and-stick with the following color code: oxygen (red), carbon (cyan), nitrogen (blue), hydrogen (white). The underlying protein structure is represented in cartoon style in grey with asparagine residues of *N*-glycosylation sites labelled in red spheres. Glycan color code: Man (green), GlcNAc (blue).

For TcruTS two amino acids, Y119 and W312, have been proposed to bind lactose and to stabilize its position in the catalytic domain once a substrate is bound in the active site (48, 49). The relative orientation of Y119 and W312 can be described as an open or stacked conformation. Consistent with the results reported by Mitchell et al. (49), the corresponding amino acids Y211 and Y408 in TconTS1 (Fig. 6 A/B, orange circles) exhibit a high flexibility and presented an open conformation with an average distance of 2.2 nm in the absence of a substrate (Fig. 7). For comparison, these two residues stayed in much closer contact in H-TconTS1, remaining at a distance of about 1.6 nm during our MD simulations (Fig. 7A).

**Figure 7:**
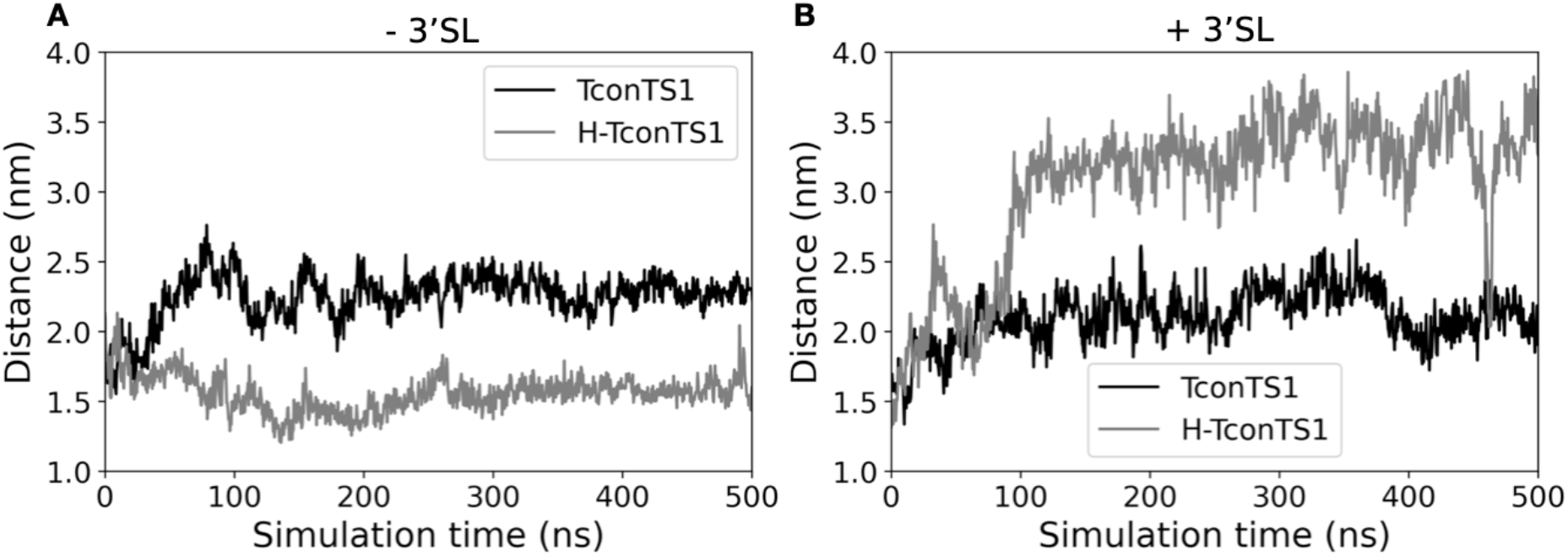
Altered movement of lactose holder amino acids in EndoHf treated H-TconTS1 compared to untreated TconTS1. Distance in nm between the Cα of Y211 and Cα of Y408 for TconTS1 (black) and H-TconTS1 (grey) measured over time in ns. A) Simulations without substrate. B) Simulations with substrate.

#### 4.3 Dynamics of conserved amino acids in the catalytic domain of substrate-bound TconTS1

As a next step, MD simulations of TconTS1 and H-TconTS1 were performed in complex with the substrate 3’SL. 3’SL was positioned in the binding site of TconTS1 in alignment with the crystal structure of the TcruTS/3’SL complex (PDB entry 1S0I). In the starting structure, 3’SL is bound at the acceptor substrate binding site between Y211 and Y408, and in close contact to both D150 and the well-conserved arginines R339, R410 (see Fig. 8A/B for TconTS1).

**Figure 8.**
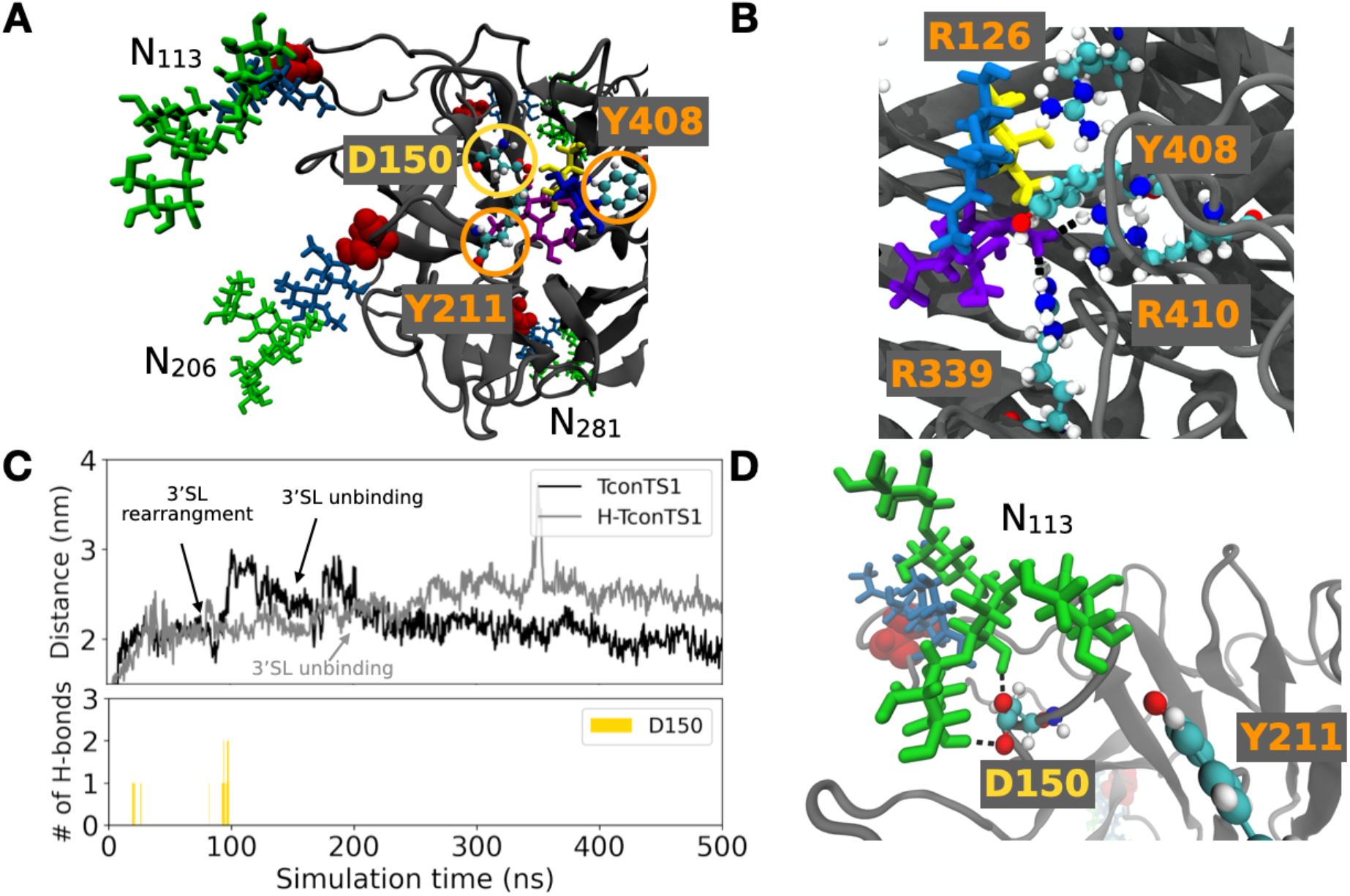
MD simulations of TconTS1 with 3’SL bound to the active center revealed protein-glycan interactions. A) Starting structure. B) Snapshot of 3’SL in the binding pocket, forming hydrogen bonds (black dashed lines) to conserved arginine residues. C) Distance between the center of D150 and the center of R410 for TconTS1 (black) and H-TconTS1 (grey). Number of hydrogen bonds between D150 and glycan N113 for TconTS1 over the course of the simulation. D) D150 interacts with glycan N113 via hydrogen bond formation (black dashed lines) at its mannose branches (snapshot at 97 ns). Color code of the amino acids as in Fig. 6. Glycan color code: Man (green), GlcNAc/Glc (blue), Gal (yellow), Neu5Ac (violet).

As already seen in the substrate-free simulation, D150 of TconTS1 formed hydrogen bonds with mannose residues of an *N*-glycan. However, in this case it was *N*-glycan at N113, which is also structurally in close proximity to the active site, and not the *N*-glycan at N206 (Fig. 8D). This interaction was observed after 20 ns, when D150 moved slightly off the catalytic site, becoming more accessible for interactions with the *N*-glycan (Fig. 8C). Following a structural rearrangement of 3’SL within the binding site after around 70 ns, D150 again interacted with *N*-glycan N113 and was dragged out of the binding site (Fig. 8C). In striking contrast, amino acids in H-TconTS1 known to be essential for the direct catalytic activity did not experience any interactions with residual GlcNAc residues, remaining after EndoH_f_ treatment.

In the presence of a substrate, different motion of Y211 and Y408 was observed with respect to the substrate-free protein. Namely, the distance between the two amino acids remained at about 2 nm in TconTS1 but increased from about 1.5. nm to more than 3 nm in H-TconTS1 in the presence of 2,3-SL (Fig. 7B).

## Discussion

In this study, we have focused on elucidating the potential influence of *N*-glycosylation on enzyme activity and stability of TconTS1 from the animal pathogen *T. congolense*. Therefore, recombinant TconTS1 was expressed in CHO Lec1 cells (31, 40). This *N*-glycosylation deficient cell line mainly yields *N*-glycans of the composition Man_5_GlcNAc_2_ due to a mutation in the *N*-acetylglucosaminyltransferase 1 (GnT1) gene (41, 50). This special type of *N*-glycosylation mimics that reported for African trypanosomes (25–30) and thus represents a suitable *N*-glycosylation model system for this study. EndoH_f_ treatment was employed for specific removal of *N*-glycans to produce a hypoglycosylated TconTS1 enzyme and study the effect on structure-function relationships. Point mutations of single *N*-glycosylation sites have not been considered in this work, as a correct folding and function of TconTS1 was important to ensure comparable enzyme activities. Co-translational *N*-glycosylation is known to influence protein folding. Our numerous experiments on site-directed glycosylation knockout using myelin associated glycoprotein (MAG; Siglec-4, unpublished data) yielded largely misfolded proteins with loss of function, whereas after enzymatic deglycosylation of the wild-type protein the function was retained. Thus, the approach of site-directed glycosylation knockout also has its limitations. Furthermore, for TconTS the orders of magnitude lower specific activity of enzyme expressed in bacteria relative to that expressed by CHO-Lec1 cells provide evidence for misfolding of the enzyme as a consequence of absence of *N*-glycans (34). Since TconTS1 exhibits nine putative *N-*glycosylation sites, a first assessment of which sites might be more important than others for enzyme function was desirable and aimed in this study.

Using a qualitative MALDI-TOF MS analyses we were able to demonstrate that at least seven out of nine potential *N*-glycosylation sites in TconTS1 are conjugated with high-mannose type *N*-glycans, of which five are located in the catalytic domain. A heterogeneous *N*-glycosylation pattern at some sites was observed especially at N206 as also higher mannose structures (Man_6-8_GlcNAc_2_) and potentially also fucosylated *N*-glycans (Man_5_GlcNAc_2_Fuc) were detected in several, but not all, mass spectra. This heterogeneity might be explained by the lack of accessibility of the glycan for glycosidases, since *N*-glycan precursors are trimmed from Man_8_GlcNAc_2_ to Man_5_GlcNAc_2_ in the Golgi (51). This hypothesis is supported by our observation that EndoH_f_ was not able to remove all *N*-glycans from TconTS1 even after 16 h of incubation. Additionally, our MD simulations revealed protein-glycan interactions between D150 and *N*-glycans attached to N113 and N206 by hydrogen bond formation. It was previously shown by biochemical *in vitro* assays and computational studies that such interactions decrease glycan accessibility, which might interfere with glycan trimming as well as with EndoH_f_-mediated removal of *N*-glycans (52). The potential presence of fucosylated *N*-glycans such as Man_5_GlcNAc_2_Fuc is a result of fucosyltransferase FUT8 activity, which was reported to act on Man_5_GlcNAc_2_ glycans (41, 53, 54). The detection of the sites N113, N206, N240 and N693 in either a glycosylated or non-glycosylated state is another form of heterogeneity already reported for other proteins (55).

The enzymatic removal of high-mannose type *N*-glycans did not disrupt the secondary structure of TconTS1. First, circular dichroism spectra of TconTS1 and H-TconTS1 did not reveal any detectable differences in secondary structure elements. Second, a similar midpoint of unfolding transition was determined for both enzymes in temperature-ramping experiments, suggesting no differences in stability. Similar results were found in single-site *N*-glycosylation mutants of the IBV spike protein by circular dichroism experiments (4). The observed thermal stability of TconTS1 might be explained by the high ß-sheet content of the protein as well as by an extended interface between catalytic and lectin-like domain stabilized by salt bridges and a well-structured hydrogen bond network, making unfolding rather unlikely (34). Although *N*-glycans do not seem to influence the stability of TconTS1 after successful expression, glycans are known to be required for proper protein folding (56). Whether or not TconTS1 requires *N*-glycosylation for accurate protein folding cannot be excluded by our experiments, and the role of *N*-glycans in TconTS folding still needs to be investigated. However, as discussed above our findings on enzymatic activity of bacterially expressed TconTS provide evidence for glycosylation-dependent misfolding of the enzymes (34). Further thermodynamic details of the denaturation processes for the two enzyme forms could be investigated in future works with methods complementary to CD, such as calorimetry.

A clear influence of *N*-glycosylation on enzymatic activity, however, was observed. Lactose affinity of H-TconTS1 was decreased by a factor of 1.6 to 5 relative to TconTS1. However, the V_max_ value was the same for both enzymes, which corroborates the structural integrity of H-TconTS1. Differences between previously reported V_max_ values for TconTS1 (4.1 ± 0.1 µmol 3’SL/(min x mg enzyme) (31) are most likely due to different enzyme preparations. It could be further argued that the glycosylated TconTS1 control in this study was incubated for 16 h at 37°C before the assays, equally to the EndoHf treated enzyme (see Methods) and that this has caused a lower V_max_. However, this is unlikely, since previous unpublished work in our laboratory has shown that prolonged incubation (even for weeks) at 37°C does not lead to loss of activity (35). Furthermore, the CD experiments of this study provided evidence for a good heat stability of TconTS1.

Haynes et al. (20) studied the influence of *N*-glycosylation on TvivTS1 and did not observe an effect on enzyme activity. The same applies to investigations of a mutated variant of TranSA, which expresses TS activity (21). However, these studies did not determine the K_M_ values for the substrates used in enzyme reactions. Another study of TranSA, in which the sialidase activity was investigated, did not observe strong effects on K_M_ when recombinant proteins were expressed in *Escherichia* (*E.*) *coli* and compared with the native enzymes isolated from trypanosomes (38). However, sialidase activity of TS is determined in the absence of an Sia acceptor substrate such as lactose. For TcruTS, enzymes expressed in *E. coli* still showed transfer activity although to a lesser extent than observed for the native protein, which might be a result of the absence of *N*-glycans and/or incorrect protein folding (36).

Molecular insight into a putative effect of *N*-glycosylation of TconTS1 at the atomic level were obtained in a series of MD simulations. The *N*-glycans, especially in TconTS1 catalytic domain, were observed to form a highly dynamical ‘shield’ enclosing the enzyme, while leaving the entrance to the catalytic center open for substrate binding.

In several simulations of TconTS1, both in the absence and presence of bound 3’SL, D150 forms either intermittent or stable hydrogen-bond interactions with *N*-glycans at positions N206 or N113. These data suggest that these *N*-glycans are involved in positioning of D150 and influence the potential of this aspartate to act as a proton donor in the enzymatic transfer of Sia, a crucial contribution to substrate conversion (16, 47).

Interestingly, these are conserved *N*-glycosylation sites among TS or SA from different species, despite variations in amino acid sequences and distributions of other *N*-glycosylation sites. In particular, N206 is conserved in all TS variants of *T. congolense* as already described by Waespy *et al*. (24) but also in enzymes from *T. cruzi*, *T. rangeli, T. brucei* and *T. vivax* as revealed by sequence alignment (Fig. S3). Additionally, N113 is also conserved in TconTS, TvivTS and TranSA. Along this line, we propose that there might be a common mechanism of TS activity mediated by *N*-glycan interactions with amino acids of the active site, in particular with D150 or its equivalents in other TS. MD simulations showed that through this interaction, which was only observed for glycosylated TconTS1, D150 was pulled out of the active site. Also the arrangement of Y211 and Y408, both expected to participate in binding of the acceptor lactose appears to be influenced by the presence of *N*-glycans. Whereas in the TconTS1 model the distance between these amino acids (about 2 nm) was not changed in the presence of the donor substrate 3’SL, in the corresponding H-TconTS1 model the two amino acids Y211 and Y408 increased from a close proximity (1.5. nm) to twice the distance (above 3 nm) if 3’SL was present in the active site.

Interaction of the *N*-glycan at position N113 with D150 was also observed in MD simulations of TconTS1 in complex with 3’SL, although more intermittent. Interestingly, the substrate was tightly bound between Y211 and Y408 in TconTS1, but not in H-TconTS1.

In summary, these observations are in agreement with the notion the *N*-glycans modulate the fine tuning of critical amino acid side chains’ arrangement at the catalytic site of TconTS1, which might facilitate the initial binding of substrates and lead to a higher substrate-binding affinity in line with our kinetic data.

It has to be noted that the heterogeneity of *N*-glycosylation at different (Table 1), was not considered in our MD simulations at this stage. Additionally, MALDI-TOF MS data suggest that position N113 and N206 are not always occupied by *N*-glycans (Table 1) but were also found to be non-glycosylated. If only one of these sites is lacking the *N*-glycan modification, the remaining one could act as a stand-in. In case both sites are non-glycosylated, such a TconTS1 form might show a similar enzyme activity as that detected for H-TconTS1 in this study. To investigate the influence of individual N-glycans or their specific structures on the catalytic activity in detail, is an elaborate task going beyond this study.

Due to the nature of MD simulations, being limited in the number of atoms to be included, only one protein was simulated at a time, either fully *N*-glycosylated at all positions experimentally identified in this study or deglycosylated with residual GlcNAc, the structure expected following complete EndoH_f_ treatment. Since experiments such as enzyme assays always represent a cumulative average of possible protein structures present in the sample, differences in the glycosylation patterns between TconTS1 preparations is very likely, but difficult to control. Thus, the experimental data are likely to reflect situations between fully glycosylated or completely deglycosylated enzymes (with residual GlcNAc), for which our MD simulations can only provide possible mechanisms at this stage. This especially applies to hypoglycosylated samples, which were shown to be only partially deglycosylated, mainly at position N206. Therefore, the kinetic effects may even be more pronounced if the glycans at N113 and N206 would be completely absent. Site-directed mutagenesis at these sites would provide such proteins, but it remains open, whether these mutants would be folded as well as the fully glycosylated TconTS1.

In conclusion, our results indicate the importance of the flexibility of the *N*-glycan shield and its potential for interactions with amino acid residues located in the catalytic site, which have not been addressed in previous studies. The short time scale and lack of *N*-glycans in previously performed MD simulations probably prevented the exploration of the D150 structural shift (48, 49, 57, 58). Detailed mechanisms underlying *N*-glycan-dependent modulation of enzymatic activities remain to be explored by advanced techniques enabling prediction of binding free energies and including hybrid quantum mechanics/molecular mechanics approaches, as well as structural NMR data, which have been beyond the scope of this study.

Our data further underline the general relevance of MD simulations, since differences in the dynamical behavior of catalytic amino acids are not always detectable in wet-lab experiments. In particular, the D150 shift could not be detected in our circular dichroism experiments, because D150 is part of a turn/coil structural motif, which does not change its secondary structural composition irrespective of the location of D150 (Fig. S8).

In summary, our study demonstrates that *N*-glycosylation can enhance substrate affinity in TS, putatively via interactions of *N*-glycans with amino acids associated with substrate binding and the catalytic process. Glycan-protein interactions are often observed, and have been suggested to regulate the conformation of proteins and their ability to bind to substrates such as collagen (59). However, to the best of our knowledge, this is the first time that interactions of the *N*-glycan shield with amino acids of the catalytically active site of an enzyme were observed to modulate enzymatic activity. As *N*-glycosylation sites are partially conserved among the trans-sialidase enzyme family, the here-observed protein-glycan interactions may also occur in TS of other trypanosome species.

## Experimental procedures

Chemicals and reagents used in this study were of cell-culture and analytical grade and were purchased from Sigma-Aldrich/Merck KGaA (Darmstadt, Germany) or Carl Roth (Karlsruhe, Germany) if not stated otherwise.

### Expression of TconTS1 in CHO Lec1 cells

Recombinant TconTS1 was produced in CHO Lec1 cells as already described (31). In brief, a modified pDEF vector was used for stable transfection comprising a transin sequence for protein secretion followed by the TconTS1-encoding sequence excluding signal peptide and GPI anchor sequence, a C3 cleavage site and a C-terminal SNAP and *Strep*-tag. Monoclonal cells were grown in serum-free CHO medium (Bio&SELL, Feucht, Germany) or Excell medium supplemented with 50 μg/mL gentamicin sulphate (Lonza™ BioWhittaker™, Walkersville, MD, USA) at 37°C and 5% CO_2_.

### Purification of TconTS1 from cell culture supernatant

Cell culture supernatant comprising the recombinant protein was harvested every second day and stabilized with 10 mM Tris/HCl pH 8.0, 10 mM EDTA, 10 mM ascorbic acid and 0.02% sodium azide. Ultracentrifugation was performed at 7,800 rcf for 15 min followed by 40,000 rcf for 45 min at 4°C. The clear supernatant was microfiltered (0.22 µm, PES) and concentrated to 50 mL using a Sartorius Vivacell 250 PES Centrifugal Concentrator (Sartorius, Göttingen, Germany) with a MWCO of 100 kDa and a pressure of 4 bar. Buffer was exchanged five times with 200 mL of 100 mM Tris/HCl pH 8.0, 150 mM NaCl, 1 mM EDTA and concentrated to a final volume of 10 mL. The concentrate was centrifuged at 21,000 rcf for 30 min and recombinant protein was purified from the supernatant with *Strep*-Tactin sepharose (IBA, Göttingen, Germany) according to the manufacturer’s protocol. Elution fractions containing TconTS1 were pooled, and buffer was exchanged four times with 5 mL of 10 mM potassium phosphate buffer pH 7.4 using a Vivaspin 6 Centrifugal Concentrator (Sartorius) at 2,000 rcf and 4°C for 20 min. Protein samples were stored at 4°C. TconTS1 concentrations were determined with a Pierce BCA Protein-Assay kit (Thermo Fisher Scientific, Schwerte, Germany) using BSA as standard.

### EndoHf treatment of TconTS1

2 mg of TconTS1 were incubated with 40,000 Units EndoH_f_ (New England Biolabs, Frankfurt am Main, Germany) for 16 h at 37°C in 2.0 mL 10 mM phosphate buffer pH 7.4. The EndoH_f_ treated enzyme is termed H-TconTS1. A TconTS1 control without EndoH_f_ addition was performed in parallel (TconTS1). The H-TconTS1 sample was purified again using *Strep*-Tactin sepharose to remove free glycans and EndoH_f_ and buffer was exchanged to 10 mM phosphate buffer pH 7.4 as already described.

### SDS-PAGE, western blot and ConA lectin blot analysis

Protein samples were separated via SDS-PAGE and either stained with PageBlue Protein Staining Solution (Thermo Fisher Scientific) or used for western blot or ConA lectin blot analysis. For western blots, a polyclonal rabbit anti-*Strep* (IBA) and a polyclonal, peroxidase-conjugated donkey anti-rabbit antibody (Jackson ImmunoResearch, Cambridgeshire, United Kingdom) were used as primary and secondary antibody, respectively. For ConA lectin blots, high-mannose *N*-glycosylated proteins were detected employing ConA-biotin (Galab, Hamburg, Germany) and the VECTASTAIN ABC-HRP Kit (Vector Laboratories, Burlingame, CA, United States).

### MALDI-TOF MS analysis of TconTS1 *N*-glycosylation sites

In-solution digestion of glycosylated and hypoglycosylated TconTS1 was performed with trypsin and chymotrypsin (Promega, Walldorf, Germany) to analyze peptides with MALDI-TOF MS. Protein solutions with a final concentration of 0.1 mg/mL were prepared in 50 mM NH_4_HCO_3_ buffer pH 7.8. Cysteine bonds were reduced with 1.4 mM DTT for 30 min at 50°C and the protein was denatured for 10 min at 95°C. After a short cooling step on ice, cysteine residues were alkylated with IAA at a final concentration of 3.2 mM for 30 min at 37°C and protected from light. Protein samples were digested with trypsin or chymotrypsin overnight at 37°C at protein:protease ratios of 25-100:1. Two negative controls were prepared accordingly, either lacking the protein TconTS1 (N1) or the protease (N2).

Samples were spotted directly on the target (Bruker Daltonics, Bremen, Germany) or glycopeptides were further purified with ConA beads as described under “Glycopeptide purification of protease-digested TconTS1 with ConA beads”. For spotting, 1 µL protein solution was directly mixed with 1 µL HCCA matrix solution (40 mM in 50% acetonitrile/0.1% TFA) on the target. Samples were measured in the positive reflector mode using the MALDI-TOF autoflex^TM^ speed (Bruker Daltonics) which was calibrated with the peptide calibration standard II (Bruker Daltonics). Detailed instrument settings can be found in Table S1. The peak assignment procedure and validation criteria are summarized in SI section S3.

### Glycopeptide purification of protease-digested TconTS1 using ConA beads

To concentrate glycopeptides from protease-digested TconTS1, samples were incubated with AffiSep® ConA adsorbent (Galab). The adsorbent was equilibrated in 1x binding buffer (50 mM Tris/HCl pH 7.4, 150 mM NaCl, 1 mM CaCl_2_, 1 mM MgCl_2_, 1 mM MnCl_2_) and was added to trypsin-digested and heat-inactivated protein samples in a ratio of 1µL adsorbent/2.5 µg protein). 5x binding buffer of appropriate volumes was supplied to samples to yield a final concentration of 1x. Samples were rotated overnight at 8 rpm and 4°C and centrifuged for 30 s at 130 rcf. The supernatant was discarded, and the adsorbent was washed thrice with 50 mM NH_4_HCO_3_ buffer (pH 7.8) in binding buffer and at the same centrifugation conditions. After the last centrifugation step the adsorbent was resuspended in 10-50 μL of NH_4_HCO_3_ buffer (pH 7.8). Elution of glycopeptides was performed at 95°C for 10 min. Samples were centrifuged at 4,300 rcf for 30 s. The supernatant was used for MALDI-TOF MS analysis. Negative controls were prepared in parallel with identical components apart from the trypsin-digested TconTS1. Peaks resulting from negative controls were excluded for data analysis.

### TS activity assay

TS activity assays were executed as previously described (31, 35). Fetuin and lactose served as Sia donor and acceptor, respectively. Reactions were carried out in 50 µL of 10 mM potassium phosphate buffer, pH 7.4 containing 100 µg dialyzed fetuin (corresponding to 600 µM bound Neu5Ac), varying concentrations of lactose (0.01-5 mM) and 50 ng hypoglycosylated TconTS1 or the corresponding control. Samples were incubated at 37°C for 30 min and the reaction was terminated with 200 µL ice-cold acetone. After protein precipitation overnight at -20°C, samples were centrifuged (20000 rcf, 30 min, 4°C), lyophilized and resuspended in 125 µL water. Transfer activity was measured as 3’SL implementing the HPAEC-PAD system ICS-5000+ (Dionex/Thermo Fisher Scientific). 25 µL sample were applied to a CarboPac100 analytical column (250×2 mm, 8.5 µm, Thermo Fisher Scientific) equipped with a guard column (50×2 mm, Thermo Fisher Scientific). Chromatography was performed at isocratic conditions with 100 mM NaOH and 100 mM NaOAc for 12 min followed by a wash step with 100 mM NaOH and 500 mM NaOAc for 5 min and an equilibration step for 8 min to previous conditions. Production of 3’SL was quantified with a purchased 3’SL standard (Carbosynth, Compton, United Kingdom). Data acquisition and evaluation was performed with the Dionex software Chromeleon 7.2 SR5. Parameters of the Michaelis-Menten equation, K_M_ and V_max_, were calculated with the curve fit model of SigmaPlot11.

### Circular dichroism experiments

Circular dichroism experiments were performed with purified recombinant TconTS1 in its glycosylated and hypoglycosylated form dissolved in 10mM phosphate buffer pH 7.4. The Applied Photophysics Chirascan spectrometer (Applied Photophysics Limited, Leatherhead, UK) with the Pro-Data Chirascan software (v.4.2.22) was used to record and evaluate circular dichroism spectra. For each sample, at least three repetitive scans were performed over a standard wavelength range of 190 to 250 nm with intervals of 1 nm. Throughout the experiments, Suprasil quartz cells (Hellma UK Ltd.) were used with a pathlength of 0.2 mm. Baseline scans were performed with 10mM phosphate buffer (pH 7.4) only, in the respective cuvette. Regarding data processing, the baseline was subtracted from recorded spectra and repetitive scans were averaged before a Savitsky-Golay smoothing filter with smoothing windows of three data points was applied. Spectra have been recorded in raw ellipticity (*Θ*) and were further converted to mean residue ellipticity

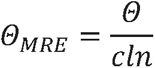

where *c* is the protein concentration, *l* the quartz cuvette pathlength and *n* the number of amino acids in the protein sequence. To be consistent with the parameters of the enzyme assay, constant-temperature measurements were performed at 35°C to check for a general influence of glycans on the protein structure.

To estimate the secondary structure components of the measured sample, circular dichroism spectra were analyzed using the BeStSel Web server (60, 61). Temperature-ramping experiments were performed following the suggestions of (62) in order to analyze protein stability, unfolding intermediates and the midpoint of the unfolding transition (melting temperature, T_M_). In detail, protein samples were heated from 20°C up to 95°C with 5°C temperature steps employing the stepped ramp mode. After 5 min of equilibration time at the respective temperature at least three spectra were recorded and averaged. The T_M_ was calculated from the fraction of protein folded at any temperature (*α*) defined as:

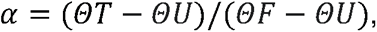

where *ΘT* is the ellipticity at any temperature, *ΘU* is the ellipticity at the unfolded state and *ΘF* at the folded state. T_M_ is defined as the temperature at which *α* = 0.5 (62) and also referred to as the melting temperature. In order to calculate *α*, we chose 195 nm as the wavelength to plot recorded *Θ_MRE_* values against the temperature. Afterwards, the calculated α values were again plotted with reference to temperature and a sigmoid curve used for fitting to obtain a precise TM value. As we did not observe a complete unfolding of TconTS1 in any temperature-ramping experiment, *ΘU* is defined as the average ellipticity of the two highest temperatures. Therefore, *ΘF* was set as the average ellipticity that was recorded for the two lowest temperatures.

### Atomistic model of glycosylated and deglycosylated TconTS1

Structures of TS from *T. congolense* have not been resolved so far by experimental techniques. Therefore, an atomistic structure of TconTS1 was modelled by the I-TASSER web server for protein structure and function predictions (63, 64) based on the recombinant sequence without the transin signal (Fig. S1 - amino acids 23-951). However, the numbering of amino acids is in correspondence with the native sequence, as depicted in Figure S1B (31). As TconTS1 was modelled with the engineered SNAP-*Strep* for consistency and better comparison with experimental data, restraints were used to achieve proper folding of this structural part. In detail, a secondary structure restraint as well as a structure template for the SNAP-*Strep* region, constructed by I-TASSER beforehand using only the SNAP-*Strep* sequence, were employed. Major templates employed by the threading algorithm were TranSA (PDB entry: 2agsA, 2A75) as well as TcruTS (PDB entry: 1ms9). Analysis and validation of homology model construction is further outlined in the SI section S5.

### Construction of the simulation cell

The freely accessible CHARMM-GUI Glycan Modeler (www.charmm-gui.org) was employed to form a disulfide bond between residues C493 and C503. For TconTS1, Man_5_GlcNAc_2_ *N*-glycans were attached at positions N45, N113, N206, N240, N281 and N693 with CHARMM-GUI (65–70) as identified with MALDI-TOF MS. Man_5_GlcNAc_2_ was chosen for all sites, as it represents the simplest and most often found *N*-glycan in CHO Lec1 cells (41). N657 was not glycosylated, although found in our MALDI experiments, because model building was already completed at the time this glycan was found. H-TconTS1 was constructed by the addition of one GlcNAc residue via a covalent linkage at positions N45, N113, N206, N240, N281 and N693. Atomistic structures of constructed models are also described in the SI section S5 (Fig S7). The simulation box was constructed with CHARMM-GUI, filling it with water molecules to obtain a distance of 15 Å between the protein and box edge. 22 K^+^ ions have been added for charge neutralization.

### Molecular dynamics simulation

All MD simulations were performed with the GROMACS 2018 version (71), using the CHARMM36m (72) force field for proteins, *N*-glycans and 3’SL (73, 74) in combination with the TIP3P water model. The leap-frog algorithm was used as an integrator and the LINCS algorithm (75) was employed to constrain bonds connected to hydrogens atoms. Temperature coupling was performed with velocity rescaling using a τ parameter of 0.1 ps (76). The verlet cut-off scheme (77) was employed for van der Waals parameters using PME and the standardized parameters suggested for CHARMM36 in the GROMACS manual version 2019.

Energy minimizations of water and ions (with restrained proteins) were performed using the steepest descent algorithm with a tolerance of 1,000 kJ mol^-1^ nm^-1^. Equilibration of water (with restrained proteins) was done in an NVT and an NPT ensemble for 1 ns, respectively. It followed the energy minimization of the proteins (with restrained water and ions) under the same conditions as before. Finally, unrestrained equilibrations were performed under NVT and NPT for 1 ns each. The production runs were performed for 500 ns in the NVT ensemble at 310.15 K, writing coordinates to file every 10 ps. A time step of 2 fs was set for all parts of the simulations if not mentioned otherwise. The systems were analyzed and visualized every 500 ps using VMD (http://www.ks.uiuc.edu/Research/vmd/), the open-source community-developed PLUMED library version 2.6 and Python version 3.7 (78–82).

### Construction of TconTS1 in complex with 3’SL

3’SL was chosen as a substrate since its composition is similar to the typical terminal branches of complex type *N*-glycans and was already used in previous enzyme assays (8). To ensure a correct binding pose of the ligand in the catalytic site, the homology-modelled TconTS1 structure was aligned to the pdb structure of TcruTS (pdb: 1s0i) in complex with 3’SL by VMD. The position of 3’SL was copied to the TconTS structure, and the ligand-protein complex was subjected to CHARMM-GUI for further processing, as described under section “Construction of the simulation cell”. Further minimization and equilibration steps were performed as described under section “Molecular Dynamics simulation”, using a timestep of 1 or 2 fs to resolve steric clashes. Production runs were performed as mentioned above.

## Data availability

The underlying data of this study, such as raw spectra of MALDI-TOF MS and circular dichroism experiments as well as structure and trajectory files of MD simulations are made available under https://doi.org/10.5281/zenodo.6102786. Plumed files used in this study are stored in the Plumed-NEST repository (21.050).

## Supporting information

This article contains supporting information: S1. Amino acid sequences; S2. Coomassie gels and blots; S3. MALDI – TOF MS; S4. Circular dichroism experiment; S5. Molecular dynamics simulations.

## Acknowledgements

The authors would like to thank Prof. Dr. Jan-Hendrik Hehemann (MARUM MPI Bridge Group Marine Glycobiology, Bremen, Germany) for the possibility of performing the HPAEC-PAD measurements in his lab and Alek Bolte for the technical support. They further thank Dr. Monika Michaelis for her assistance with circular dichroism measurements and for fruitful discussions.

## Author contributions

Conceived and designed the experiments: JR ILG LC SK MW. MALDI-TOF MS experiments: JR ILG YY, Enzyme assays: JR ILG YY NDK, Circular dichroism: ILG YY, Molecular dynamics simulation: ILG. Data analysis: JR ILG. Manuscript writing: JR ILG LC MW. Manuscript revision: JR ILG LC SK MW.

## Funding and additional information

Computational resources were provided by the North German Supercomputing Alliance (HLRN) under the project number hbb00001. This study’s financial support came from the Deutsche Forschungsgemeinschaft (DFG: https://www.dfg.de/), project grants to SK, Ke428/10-1 and Ke428/13-1). The funders had no role in study design, data collection and analysis, decision to publish, or preparation of the manuscript.

## Abbreviations

3’SL: 3’-sialyllactose
Å: Angström
ConA: concanavalin A
H-TconTS1: EndoH_f_ - treated TconTS1
*E*.: *Escherichia*
EndoH_f_: recombinant protein fusion of Endoglycosidase H and maltose binding protein
fs: femtosecond
Gal: galactose
Glc: glucose
GlcNAc: *N*-Acetylglucosamine
HPAEC-PAD: High performance anion exchange chromatography with pulsed amperometric detection
K: Kelvin
kJ: kilo Joule
MALDI-TOF: matrix-assisted laser desorption ionization time-of-flight
Man: mannose
MD: Molecular dynamics
MS: mass spectrometry
Neu5Ac: *N*acetylneuraminic acid
nm: nanometer
ns: nanosecond
ps: picosecond
P.: Pichia
SA: sialidase
Sia: sialic acid
*T*.: *Trypanosoma*
TcruTS: *Trypanosoma cruzi* trans-sialidase
TconTS: *Trypanosoma congolense* trans-sialidase
TM: melting temperature
TranSA: *Trypanosoma rangeli* sialidase
TS: trans-sialidase
TvivTS: *Trypanosoma vivax* trans-sialidase.

## Supporting Information

### S1. Amino acid sequences

A recombinant TconTS1 construct was used for protein expression as described by Koliwer-Brandl et al. (31) (Fig. S1A). In the construct, a transin signal sequence replace the native N-terminal signal peptide, and a SNAP-*Strep* tag at the C-terminus the GPI anchor sequence (Fig. S1B). This recombinant protein allows for the purification from cell media due to its secretion from mammalian cells and a facilitated detection by the *Strep*-tag. Details about construct building and expression are given in reference (31). An overview of the recombinant construct is given in Figure S2.

**Figure S1:**
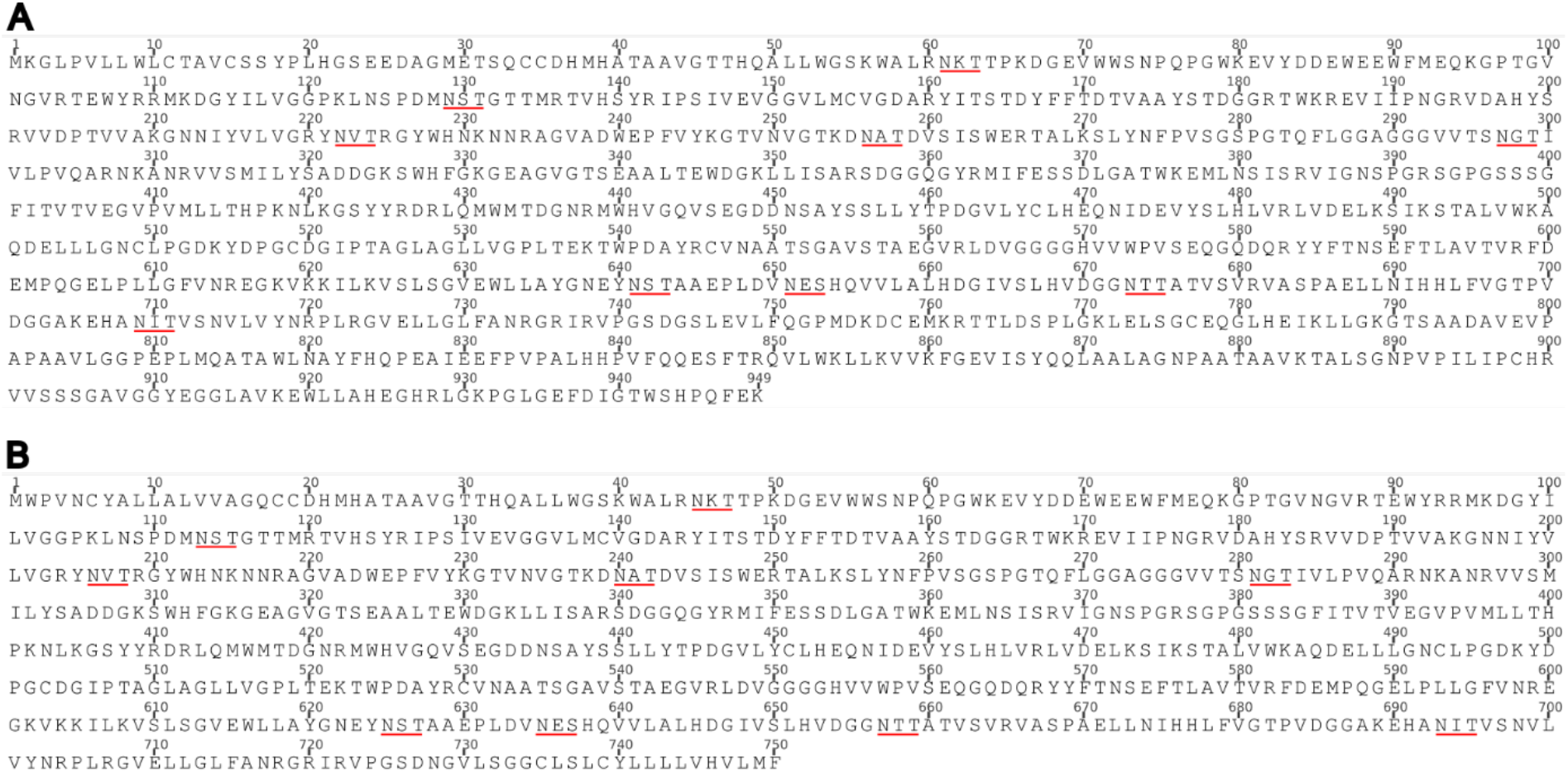
A) Amino acid sequence of recombinant TconTS1 (see Figure S2) used in this study. The signal peptide sequence of the native protein (B, GenBank ID HE583284) was replaced by a transin sequence for efficient cell secretion and the GPI anchor sequence was replaced by a SNAP- and *Stre*p-tag for protein labelling and purification. To be congruent with previous literature about TconTS1, the amino acid numbering of the native protein was used in the text. A schematic structure of the recombinant truncated protein is given in Figure S2. *N*-glycosylation sites are underlined in red.

**Figure S2:**
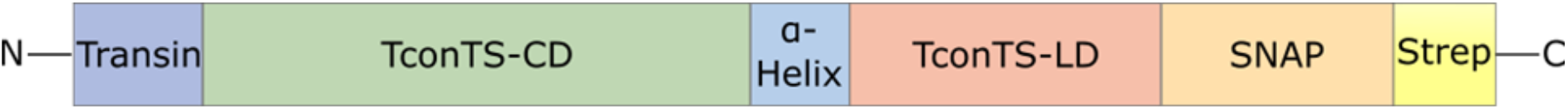
Naturally occurring TS are composed of a catalytic and lectin-like domain (CD, LD) linked by an α-helix. The recombinant protein construct of this study contained the TconTS1 coding sequence without the naturally occurring signal peptide and GPI anchor sequence and was additionally supplied with an N-terminal transin sequence for cell secretion and a C-terminal SNAP- and *Strep*-tag for protein labelling and purification, respectively.

An alignment of several TS sequences from different species (Fig. S3) shows the overall distribution of *N*-glycosylation sites marked by violet triangles. Overlaps of *N-*glycosylation positions are especially visible for the sites N113 and N206, according to the TconTS1b nomenclature labelling. These sites are indicated by red squares and are identical in up to 3-4 different TS species. Namely, N113 is conserved between TranSA, TvivTS1 and TconTS1b and N206 between TcruTS, TranSA, TbruTS and TconTS1b. This consistency of *N*-glycosylation sites might hint at conserved features, which could be important for enzyme structure or function. Overall, all TS species investigated here harbor at least two *N*-glycosylation sites.

**Figure S3:**
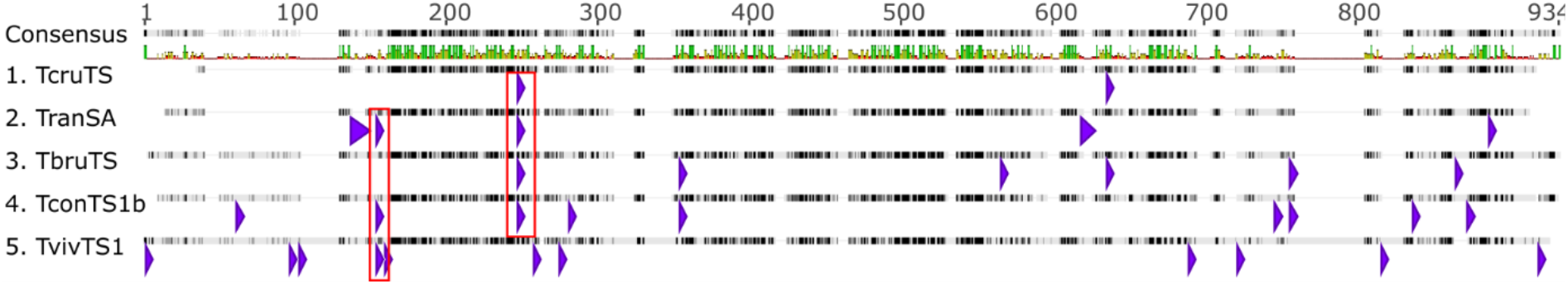
Sequence alignment of TS from *T. cruzi* (TcruTS, GenBank ID AAA66352), *T. brucei brucei* (TbruTS, GenBank ID AAG32055), *T. congolense* (TconTS1b, GenBank ID HE583284), *T. vivax* (TvivTS, GenBank ID CCD21087) and the closely related sialidase from *T. rangeli* (TranSA, GenBank ID AAC95493) was generated with the ClustalW Alignment tool of the software Geneious Pro 5.5.9 employing the BLOSUM matrix with a gap opening cost of 10 and gap penalty cost of 0.1. Conserved glycosylation sites are indicated (red box).

### S2. Coomassie gels and blots

In order to verify and assess the efficiency to deglycosylate TconTS1 using EndoH_f_, several techniques have been used including SDS-PAGE with Coomassie staining, western blotting using an anti-*Strep*-tag antibody and lectin blotting using ConA for detection of high-mannose type *N*-glycans. TconTS1 samples have always been incubated at the same temperature and for the same time as H-TconTS1, only lacking EndoH_f_.

First, EndoH_f_ treatment has been performed for 4 hours (Fig. S4A), leading to a clear shift of H-TconTS1 from ∼120 to ∼110 kDa in Coomassie and western blot analysis, indicating a mass loss due to removal of high-mannose type *N*-glycans. The ConA blot, however, indicated that complete removal of *N*-glycans was not achieved. A band was still visible at 110 kDa for H-TconTS1, although much weaker compared to TconTS1. The original and fully displayed gels of Fig. S4B also showed the presence of EndoH_f_ at ∼70 kDa in H-TconTS1, but not in TconTS1 samples.

Due to the lack of complete *N*-glycan removal after 4 hours of incubation, different incubation times of up to 48 hours or higher amounts of EndoH_f_ enzyme were tested. Complete removal of high-mannose type *N*-glycans could still not be achieved, probably due to low accessibility of the *N*-glycan for EndoH_f_ (data not shown).

In the end, we decided for an overnight incubation of 16 hours with EndoH_f_ and used these samples in all following experiments like MALDI-TOF MS, enzyme assays and circular dichroism (Fig. S4C). Also, in these blots a clear reduction of mass for H-TconTS1 bands is visible. Bands are not as clear and thin as in Figure S3A, as around 5 times more sample was loaded on all gels to detect possible degradation products. An overloading/overexposure was therefore visible. The thickness of bands might additionally result from a certain *N-*glycan heterogeneity in the samples, resulting in different migration behaviors. Original gel and blots in Fig. S4D of Fig. S4C indicate several bands in western and ConA blots at masses around 40 – 70 kDa. These bands are likely fragmentation products of TconTS1, as they could still be detected with the anti-*Strep*-Tag antibody, indicating the presence of the *Strep*-tag, and harbor *N*-glycans as detected in the ConA blot. However, the Coomassie gel in Fig. S3D clearly showed the TconTS1 band at 120 kDa and H-TconTS1 band at 110 kDa as the most prominent proteins, indicating a mostly intact protein in the sample. These results also underline the sensitivity of western and ConA blots for small amounts of protein, as the western blot showed the same signal intensity for bands around 120 kDa and around 50 kDa, although the Coomassie gel unambiguously demonstrates the different protein amount for the different bands.

**Figure S4:**
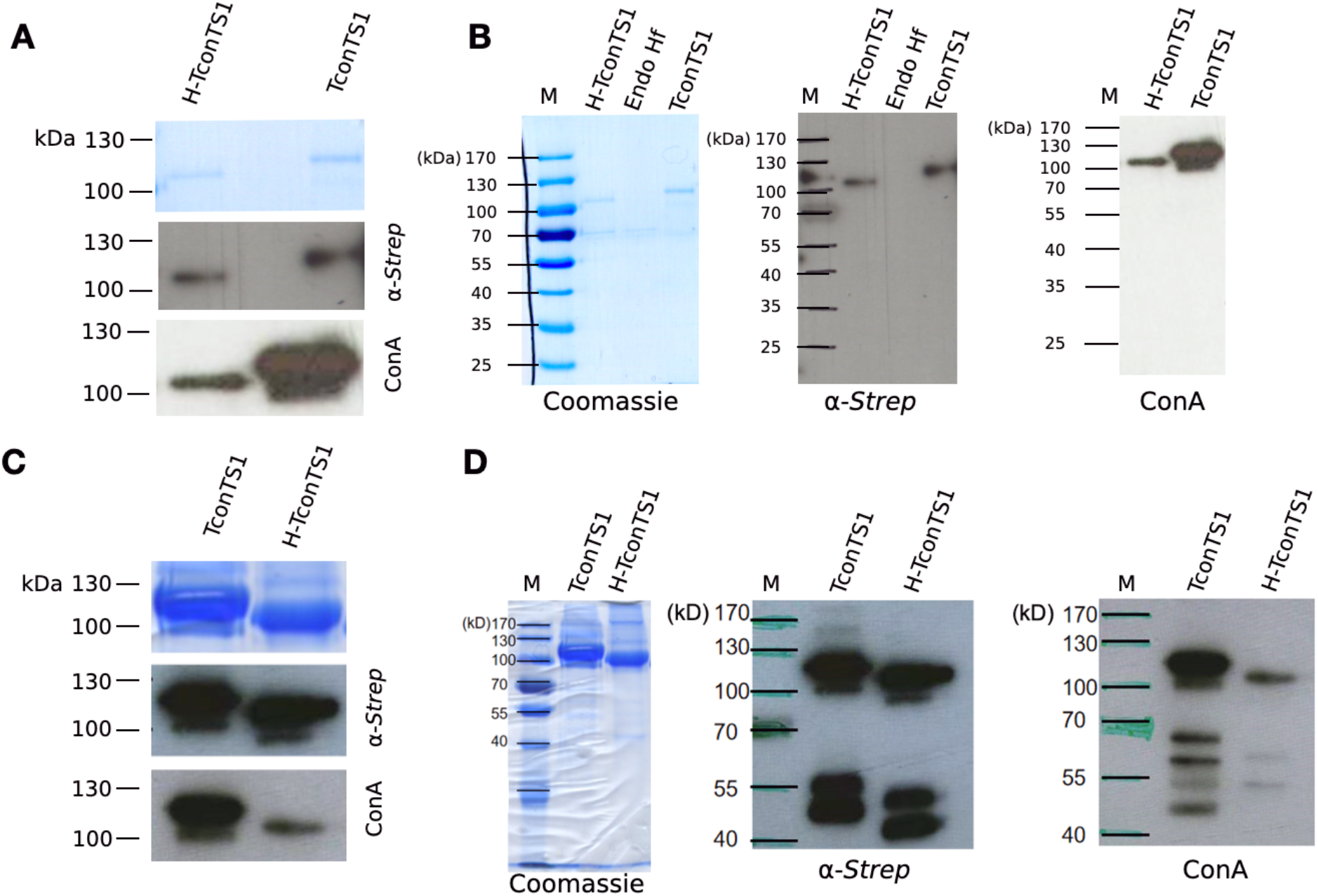
A) Coomassie staining (upper panel with 600 ng), western blot analysis using an anti-*Strep*-tag antibody (middle panel with 400 ng of protein in each slot) and ConA (lower panel with 100 ng of protein) of TconTS1 and H-TconTS1 after 4 h of EndoHf treatment. Exposure time of western blot is 5 sec and 60 sec for the ConA blot. Identical to Fig. 2A. B) Full blots used in A) with original marker bands (M). Blots were performed before purification of H-TconTS1 by *Strep*-Tactin® Sepharose affinity chromatography to remove the EndoHf enzyme. A positive control of EndoHf is loaded for the Coomassie gel and western blot. Faint bands in the Coomassie gel at 70 kDa correspond to the EndoHf enzyme. C) TconTS1 and H-TconTS1 (EndoHf-treated for 16 h) were analyzed by SDS-PAGE with subsequent Coomassie staining (upper panel with 5 µg), by western blot analysis using an anti-*Strep*-tag antibody (middle panel with 1 µg) and by lectin blotting using ConA (lower panel with 1 µg). western and ConA blots were exposed for 5 and 10 sec, respectively. D) Original blots for C) with original marker bands (M).

### S3. MALDI – TOF MS

TconTS1 harbors nine potential *N*-glycosylation sites, five in the *N*-terminal catalytic and four in the lectin-like domain. MALDI-TOF MS was employed to assess the glycosylation status of each *N*-glycosylation site.

Detailed instrument settings for this analysis are given in Table S1. Furthermore, the software Flex analysis 3.4 (Bruker Corporation, USA) was employed for data evaluation. Monoisotopic peaks were identified with Sophisticated Numerical Annotation Procedure (SNAP) algorithm and a signal to noise ratio of 6. Further settings included baseline subtraction (TopHat) and spectra smoothing (algorithm: Savitzky Golay). The online tools PeptideMass (84), FindPept (85) and GlycoMod (86) (used from 2019-2021) were used to determine (glyco)peptides as [M+H]^+^ ions and glycan structures assuming especially the presence of high-mannose type *N*-glycans as identified on proteins expressed in CHO Lec1 cells (41). Differences in m/z of 0.1 Da were accepted for identification. An exception was the analysis of TconTS1 (trypsin-digested, ConA purified) since m/z ratios of all identified glycopeptides with *N*-glycosylation sites differed by ca. 0.2 Da most likely due to the calibration. Thus, 0.2 Da was accepted for this analysis which was also the default setting of GlycoMod (86). Peaks identified in negative controls N1 and N2 were excluded from sample peak lists. In addition to glycan residues, modifications like oxidation of methionine and alkylation of cysteine were considered during peak assignment.

**Table S1:** Detailed settings of the MALDI-TOF autoflex^TM^ speed method used for identification of glycosylated and non-glycosylated peptides from TconTS1 and H-TconTS1.

| Parameter | Setting |
| --- | --- |
| Mass range | 500 – 4500/6000 |
| Smartbeam Parameter Set | Medium |
| Frequency | 500 Hz |
| Sample rate | 4 Gs/s |
| Baseline offset adjustment | 1.2 % |
| Analog offset | 51 mV |
| Ion source 1 | 19.11 kV |
| Ion source 2 | 16.91 kV |
| Lens | 8.69 |
| Reflector 1 | 20.98 kV |
| Reflector 2 | 9.44 kV |
| Pulsed ion extraction | 130 ns |
| Matrix suppression mode (up to) | Deflection (500) |
| Polarity | Positive |
| Detector gain voltage | 1700 V |

In order to analyze (H-)TconTS1 with MALDI-TOF MS, it was necessary to cut the enzyme in shorter peptide and glycopeptide fragments via protease digestion. In case an asparagine residue in the N-X-S/T motif of a certain glycopeptide is glycosylated the m/z ratio increases by exactly the mass of the conjugated *N*-glycan relative to the non-glycosylated peptide.

Three different approaches were used for peak assignment, as instrument settings did not allow for peak fragmentation and hence an unambiguous assignment of peaks. However, we believe that the combination of following experiments is suitable for cross-validation of identified peaks:

1. The usage of trypsin or chymotrypsin for generation of glycopeptides yields altered peptide profiles due to different specific protease recognition sites (Fig. S5B). The same *N*-glycosylation site can therefore be identified from different glycopeptides (different size/mass). If glycopeptides can be identified from trypsin and chymotrypsin digested samples, there is a high possibility that the assigned peak is the predicted glycopeptide. Along this line, if the mass increase is identical for glycopeptides containing the same *N*-glycosylation site when compared to the non-glycosylated peptide, this is a strong indication that this site is occupied with a defined *N*-glycan.
2. Subsequent ConA-sepharose purification of glycopeptides yielded only glycopeptides with high-mannose type *N*-glycans in the sample. As a consequence, only glycosylated fragments can be measured and identified. A further advantage of this approach is a reduced peak complexity in spectra as most peptide fragments are filtered out (Fig. S5A).
3. Further, H-TconTS1 was digested with trypsin and chymotrypsin. This step is an additional validation for the presence of *N*-glycans as EndoH_f_ cleaves the *N*-glycan tree in the chitobiose core, leaving one GlcNAc residue to the *N*-glycosylation site. These peptides fragments in H-TconTS1 samples can be detected as they exhibit a mass difference of m/z 203.08 (one HexNAc) relative to the non-glycosylated peptide. A remaining GlcNAc residue indicates the presence of a high-mannose type *N*-glycan at that site before removal by EndoH_f_ (Fig. S5C).

A detailed list of all assigned peaks with theoretical and measured masses is given in Table S2. For a better overview Table S3 (identical to Table 1 in the main text) has been constructed, separating results by the three different approaches employed for cross-validation of *N*-glycan detection and are discussed in detail in the following paragraphs.

**Figure S5:**
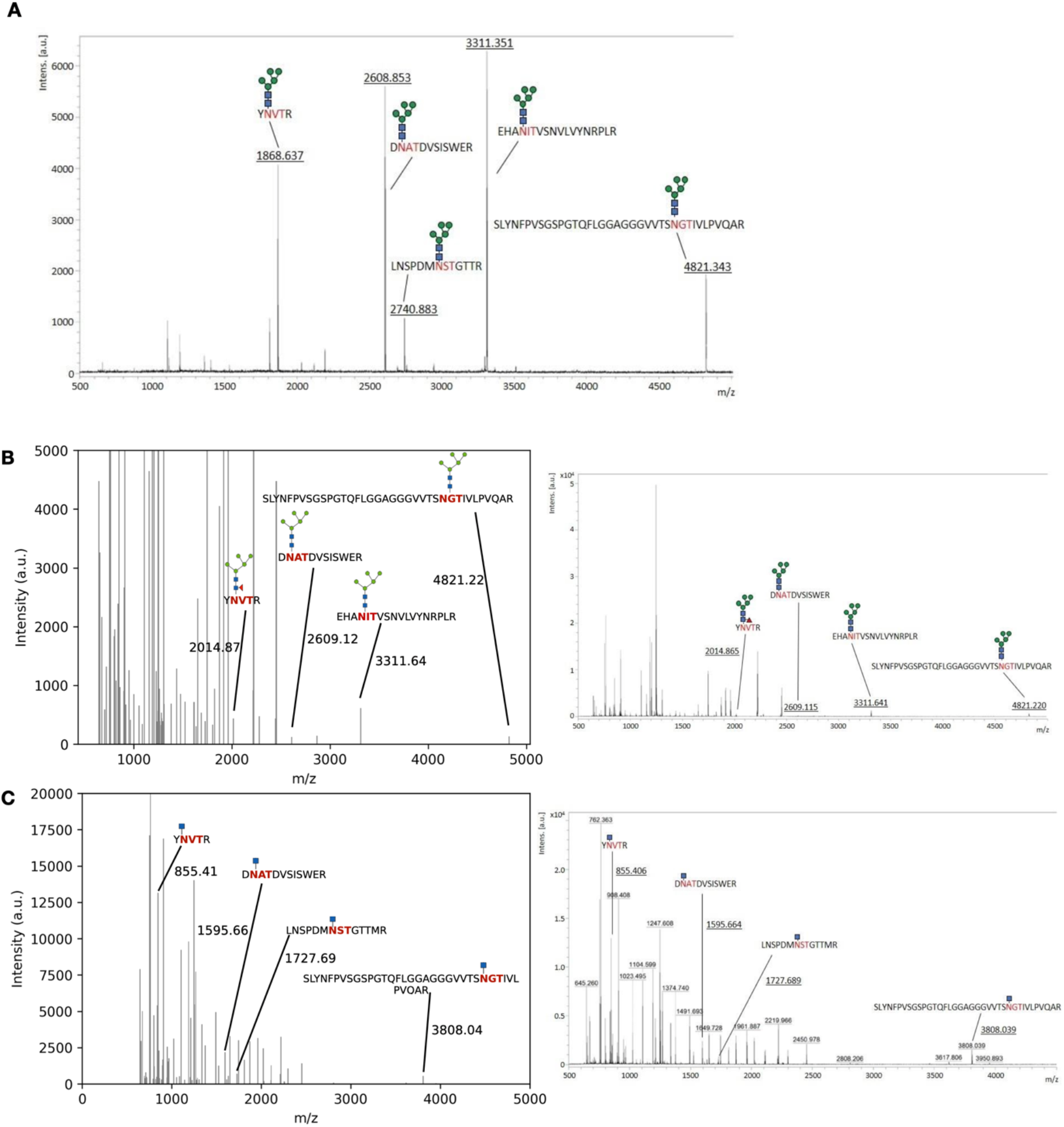
A) Original and fully unzoomed MALDI-TOF mass spectrum of trypsin-digested and ConA-purified TconTS1 from Fig. 2B. B) MALDI-TOF MS analysis of protease-digested TconTS1 identified glycosylated and non-glycosylated peptides. C) MALDI-TOF MS analysis of H-TconTS1 confirmed high-mannose type *N*-glycosylation of several glycopeptides as the EndoHf treatment only cleaves these specific glycans resulting in residual HexNAc. Peak lists from spectra were extracted and plotted with python and annotated with corresponding masses and glycopeptide fragments, respectively. They can be compared to their original, unzoomed spectra on the right (B/C). Monosaccharide symbols follow the Symbol Nomenclature for Glycans (SNFG) (83).

**Table S2:**
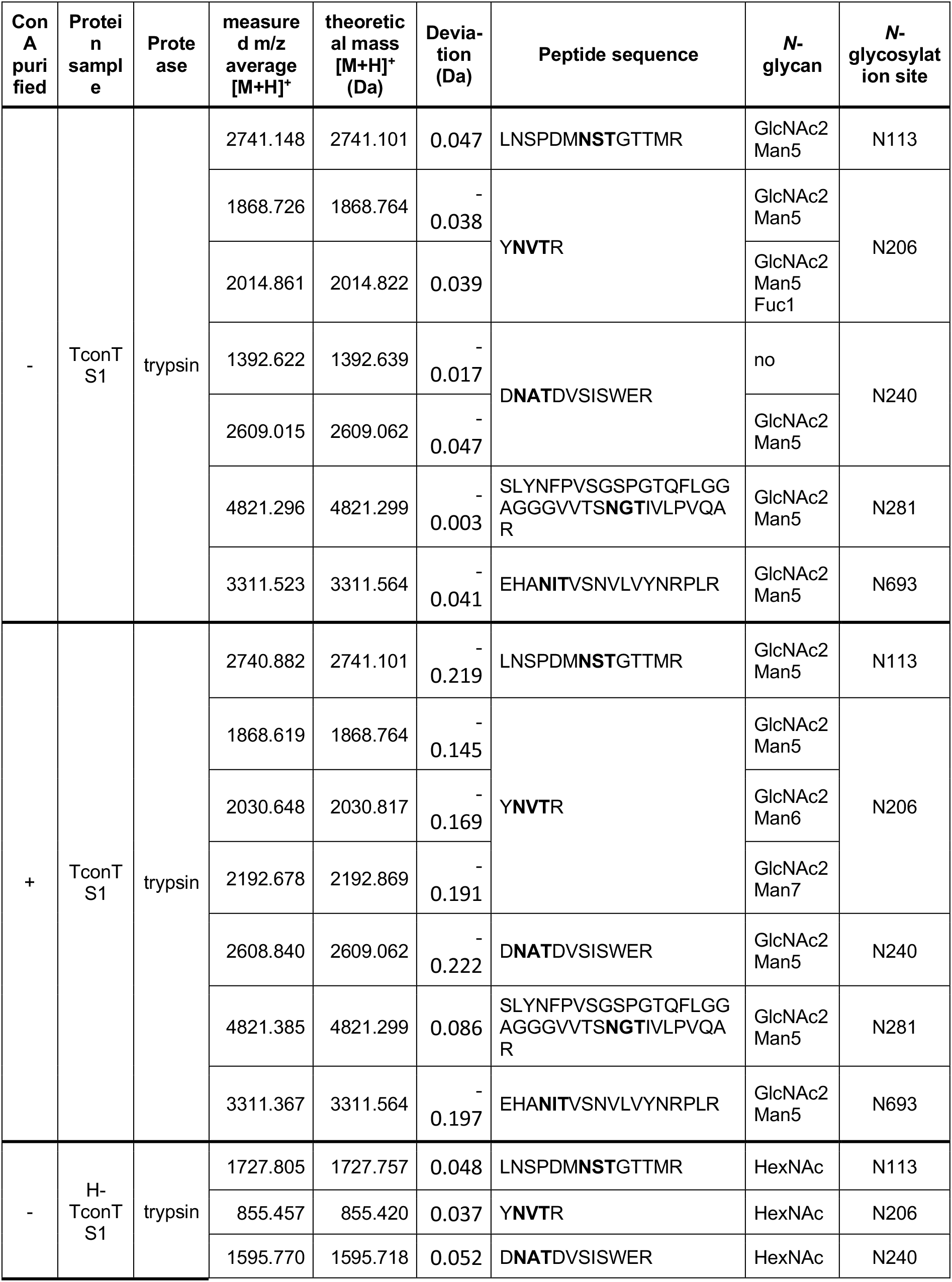
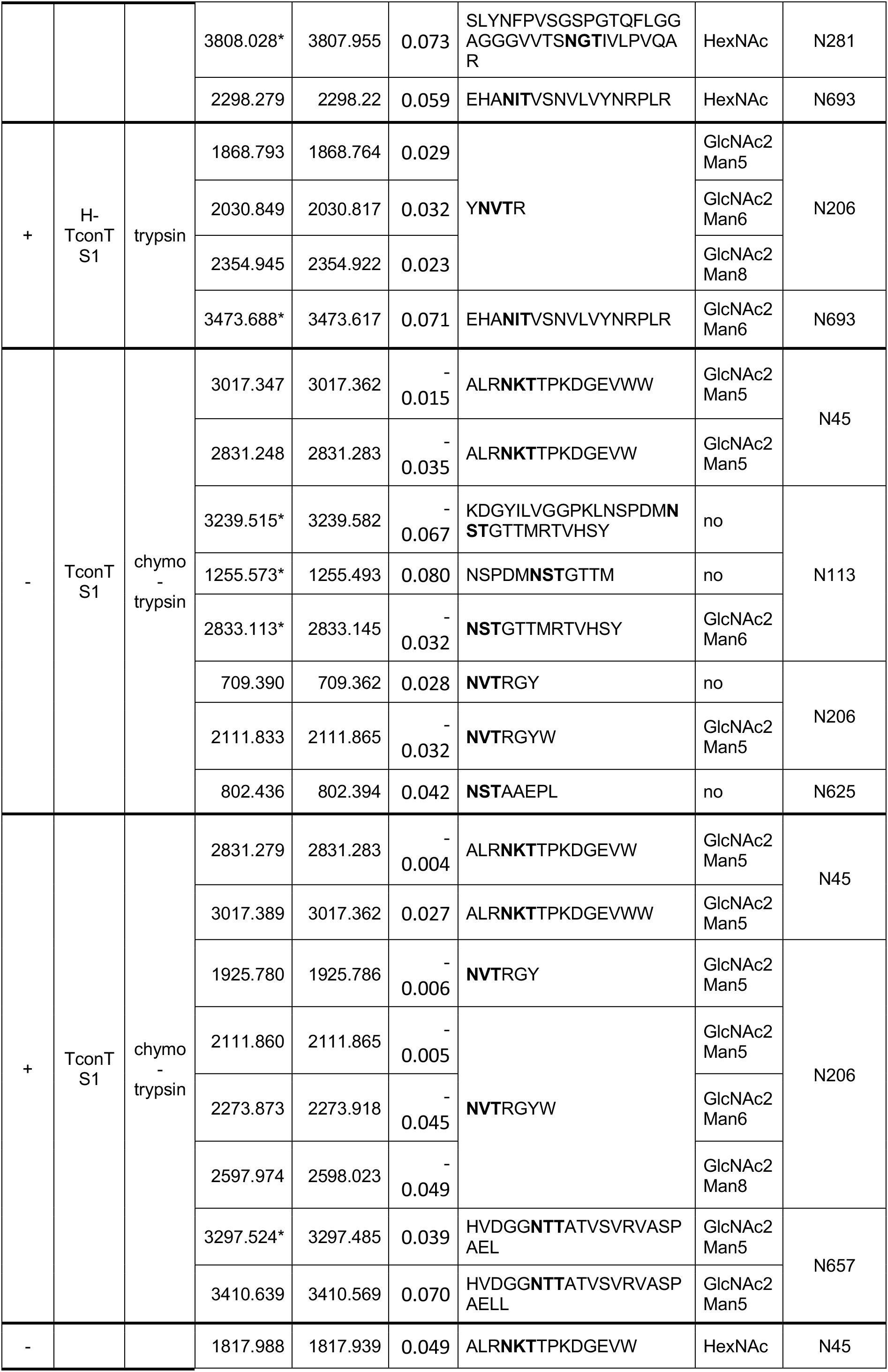
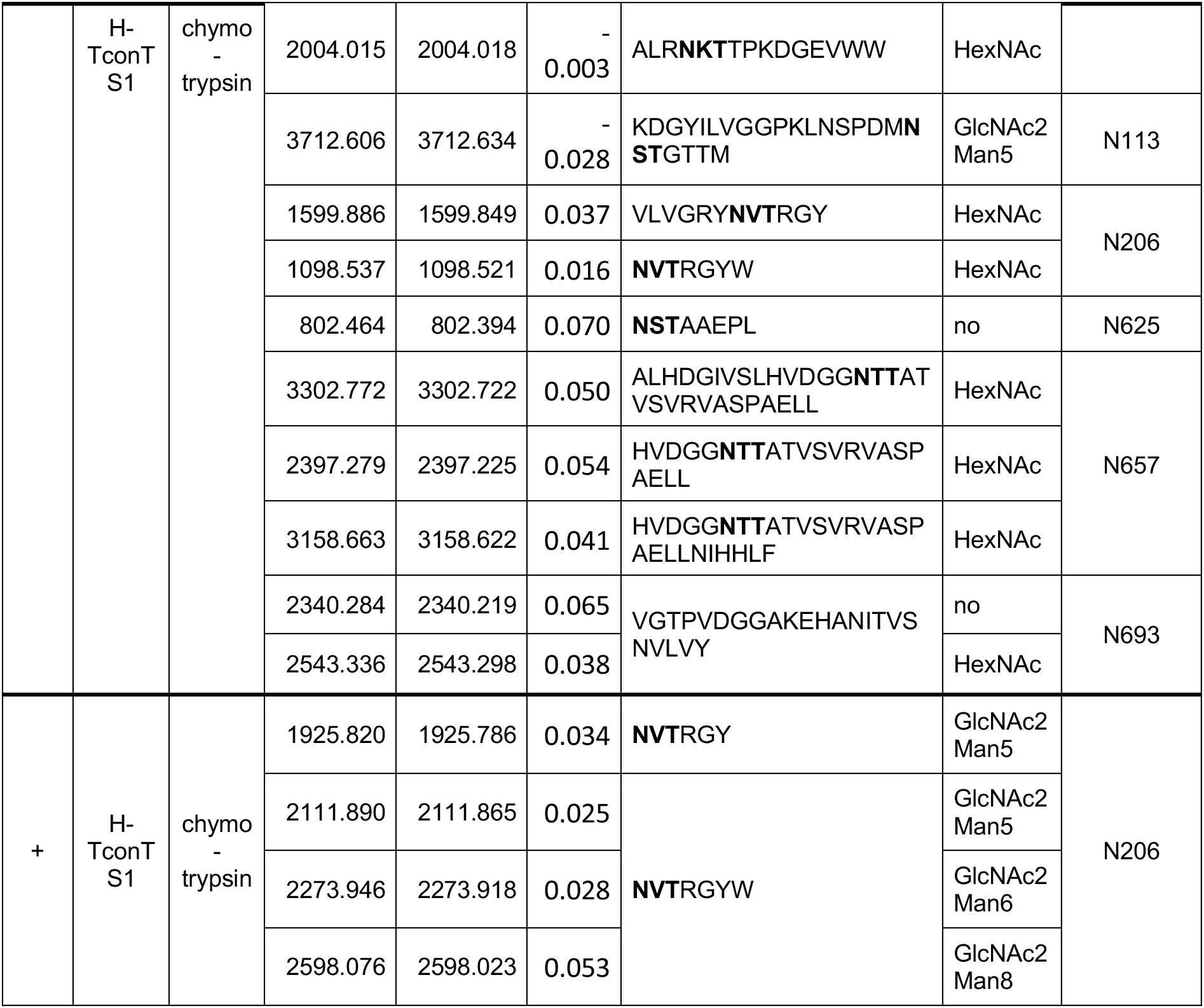
List of peptides with N-glycosylation sequences identified in MALDI-TOF mass spectra. TconTS1 and H-TconTS1 were digested with trypsin and chymotrypsin and glycopeptides were optionally purified with ConA adsorbent. Mass spectra were analyzed for the presence of peptides with the N-X-S/T recognition sequence and for the corresponding glycopeptides with high-mannose type N-glycans or HexNAc residues (H-TconTS1 only). Ions measured were only charged once ([M+H]^+^). Masses of glyco(peptide) fragments including modifications like oxidation of methionine and alkylation of cysteine could not be detected. *(Glyco)peptides only identified once during multiple analyses.

**Table S3:**
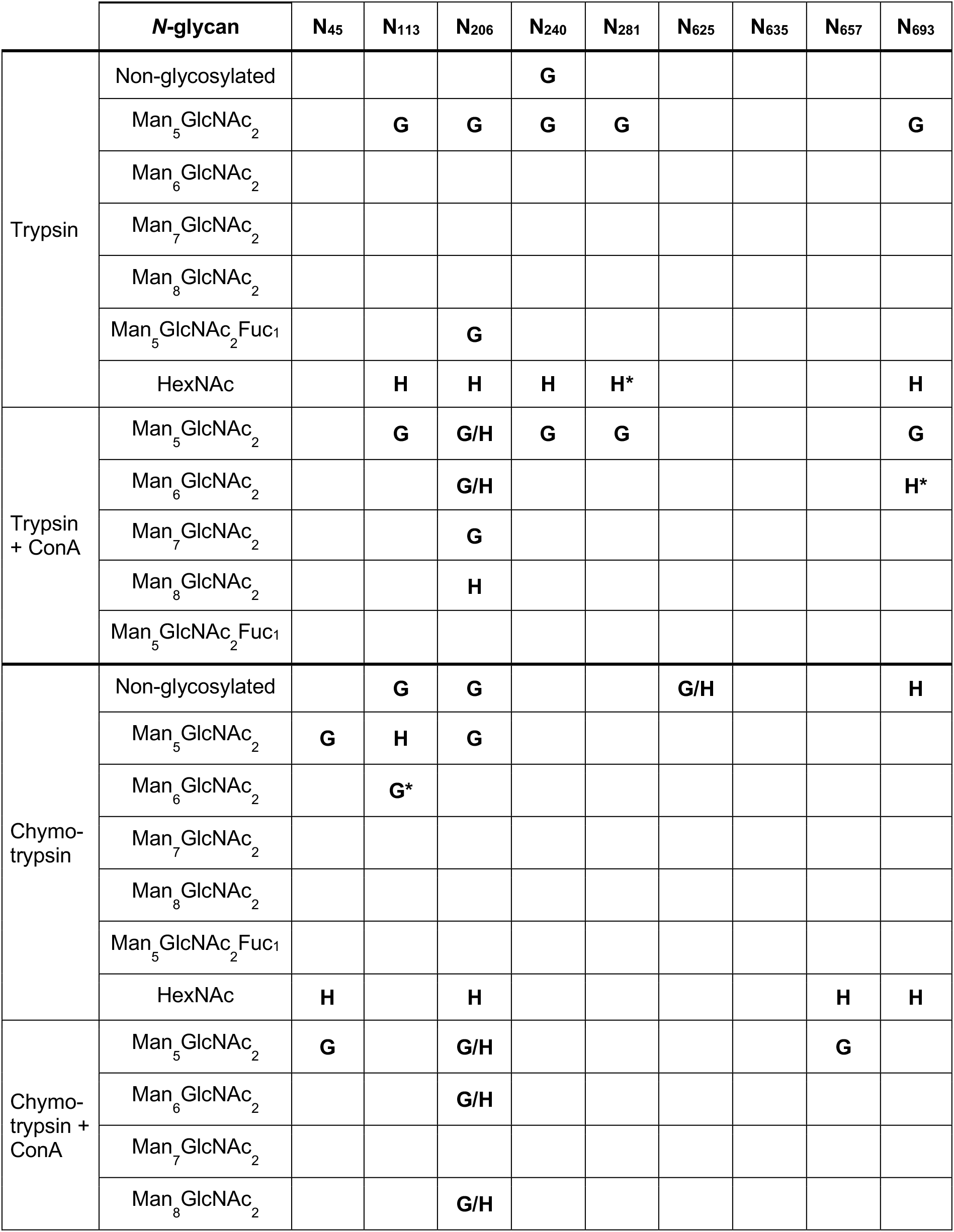
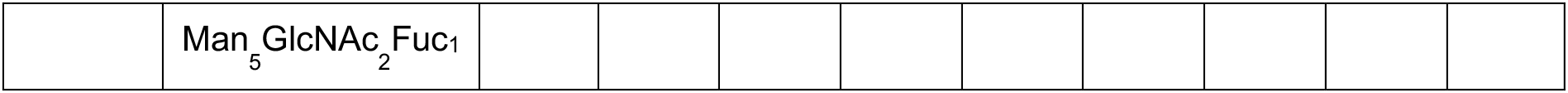
MALDI-TOF MS analyses of the *N*-glycan profile of TconTS1 (identical to Table 1 in the main text). For analysis, the untreated TconTS1 (glycosylated, G) and the EndoHf treated H-TconTS1 (H) were digested with trypsin or chymotrypsin. Spectra were analyzed for masses corresponding to glycopeptides with high-mannose type *N*-glycans and for non-glycosylated peptides with potential *N*-glycosylation sites. Spectra of H-TconTS1 were additionally analyzed for glycopeptides with HexNAc residues since a residual GlcNAc remains attached to the protein *N*-glycosylation sites after EndoHf treatment. In another approach, glycopeptides from both proteins were purified with ConA after protease digestion and spectra were analyzed for masses of peptides with high-mannose type *N*-glycans. *Glycopeptide mass was only detected once during multiple analyses.

|  | ***N*-glycan** | **N<sub>45</sub>** | **N<sub>113</sub>** | **N<sub>206</sub>** | **N<sub>240</sub>** | **N<sub>281</sub>** | **N<sub>625</sub>** | **N<sub>635</sub>** | **N<sub>657</sub>** | **N<sub>693</sub>** |
| --- | --- | --- | --- | --- | --- | --- | --- | --- | --- | --- |
| Trypsin | Non-glycosylated |  |  |  | **G** |  |  |  |  |  |
|  | Man<sub>5</sub>GlcNAc<sub>2</sub> |  | **G** | **G** | **G** | **G** |  |  |  | **G** |
|  | Man<sub>6</sub>GlcNAc<sub>2</sub> |  |  |  |  |  |  |  |  |  |
|  | Man<sub>7</sub>GlcNAc<sub>2</sub> |  |  |  |  |  |  |  |  |  |
|  | Man<sub>8</sub>GlcNAc<sub>2</sub> |  |  |  |  |  |  |  |  |  |
|  | Man<sub>5</sub>GlcNAc<sub>2</sub>Fuc<sub>1</sub> |  |  | **G** |  |  |  |  |  |  |
|  | HexNAc |  | **H** | **H** | **H** | **H\*** |  |  |  | **H** |
| Trypsin + ConA | Man<sub>5</sub>GlcNAc<sub>2</sub> |  | **G** | **G/H** | **G** | **G** |  |  |  | **G** |
|  | Man<sub>6</sub>GlcNAc<sub>2</sub> |  |  | **G/H** |  |  |  |  |  | **H\*** |
|  | Man<sub>7</sub>GlcNAc<sub>2</sub> |  |  | **G** |  |  |  |  |  |  |
|  | Man<sub>8</sub>GlcNAc<sub>2</sub> |  |  | **H** |  |  |  |  |  |  |
|  | Man<sub>5</sub>GlcNAc<sub>2</sub>Fuc<sub>1</sub> |  |  |  |  |  |  |  |  |  |
| Chymo-<br>trypsin | Non-glycosylated |  | **G** | **G** |  |  | **G/H** |  |  | **H** |
|  | Man<sub>5</sub>GlcNAc<sub>2</sub> | **G** | **H** | **G** |  |  |  |  |  |  |
|  | Man<sub>6</sub>GlcNAc<sub>2</sub> |  | **G\*** |  |  |  |  |  |  |  |
|  | Man<sub>7</sub>GlcNAc<sub>2</sub> |  |  |  |  |  |  |  |  |  |
|  | Man<sub>8</sub>GlcNAc<sub>2</sub> |  |  |  |  |  |  |  |  |  |
|  | Man<sub>5</sub>GlcNAc<sub>2</sub>Fuc<sub>1</sub> |  |  |  |  |  |  |  |  |  |
|  | HexNAc | **H** |  | **H** |  |  |  |  | **H** | **H** |
| Chymo-<br>trypsin +<br>ConA | Man<sub>5</sub>GlcNAc<sub>2</sub> | **G** |  | **G/H** |  |  |  |  | **G** |  |
|  | Man<sub>6</sub>GlcNAc<sub>2</sub> |  |  | **G/H** |  |  |  |  |  |  |
|  | Man<sub>7</sub>GlcNAc<sub>2</sub> |  |  |  |  |  |  |  |  |  |
|  | Man<sub>8</sub>GlcNAc<sub>2</sub> |  |  | **G/H** |  |  |  |  |  |  |

|  |  |
| --- | --- |
|  | Man<sub>5</sub>GlcNAc<sub>2</sub>Fuc<sub>1</sub> |

When using trypsin for protease digestion, all identified glycopeptides (N113, N206, N240, N281, N693) could also be found deglycosylated in H-TconTS1 with a residual HexNAc, indicating a reliable assignment of peaks. These results can be further underlined as the same glycopeptide fragments could be assigned in ConA sepharose purified samples, too. Glycopeptide fragments with high-mannose type *N*-glycans were also identified in H-TconTS1 samples, indicating an incomplete deglycosylation process. However, these fragments could only be identified in ConA sepharose purified samples but not in the complete trypsin digest. This indicates a low frequency of these hypoglycoslyated peptides after treatment with EndoH_f_. Identifying site N240 in a non-glycosylated and glycosylated state indicates glycosylation heterogeneity, which is possible when enzymes of the glycosylation machinery do not have temporal or spatial access to certain sites.

Looking at the chymotrypsin protease digestion, sites N45 and N206 can again be unambiguously defined in glycoproteins, harboring high-mannose type *N*-glycans in TconTS1 and one HexNAc in H-TconTS1. At N113, high-mannose type *N*-glycans were found in TconTS1 but also in H-TconTS1 samples indicating resistance to EndoH_f_ cleavage. The sites N113, N206, and N693 were also identified in a non-glycosylated state in TconTS1 or H-TconTS1 samples. N625 was exclusively found non-glycosylated and never with high-mannose type *N*-glycans or one HexNAc residue, indicating that this site might not harbor *N*-glycans at all. Chymotrypsin digestion in combination with ConA sepharose purification underlines the glycosylation status at N45, N206 and N657 as mainly Man_5_GlcNAc_2_ were identified at these sites. Additionally, site N657 was found with a residual HexNAc in simple chymotrypsin digestion of H-TconTS1, being in agreement with the Man_5_GlcNAc_2_ identified in TconTS1. The presence of high-mannose type *N-*glycan at position N206 in H-TconTS1 samples confirm the results obtained from ConA sepharose purified and trypsin-digested samples towards an incomplete deglycosylation at this site using EndoH_f_.

It seems likely that different proteases produce certain (glyco)peptides with a higher frequency and that miscleavages due to possible protease resistance of glycosylated protein at some sites might also play a role since certain peptides were neither detected in the unmodified nor in a modified status, e. g. with oxidized methionines or alkylated cysteines.

Only a few glycopeptides were identified with only one approach, such as residual high-mannose type *N*-glycans after EndoH_f_ treatment at position N113 and N693 or fucosylated Man_5_GlcNAc_2_ at N206 in TconTS1. We assume that these glycopeptides were rather rare in samples and were therefore not identified with different analyses. As these results are ambiguous, they were not regarded further in this study.

In none of these experiments we could identify a leucine-rich peptide containing the *N*-glycosylation consensus sequence N-E-S (662–664), neither in the glycosylated nor in the non-glycosylated form. The peptide was not detectable also when the mass lists were analyzed for different *N*-glycans and post-translational modifications. Among the many different proteases which are suitable for MALDI-TOF MS analysis, the leucine-rich fragment only contained cleavage sites for chymotrypsin. However, the protease cleaves at the carboxyl side of different amino acids (Y, F, W, L, M), but the cleavage efficiency is rather low after leucine and methionine. It is therefore possible that the leucine-rich fragment was not sufficiently cleaved by chymotrypsin and thus too large to be detected. Another possibility is that the fragment has been digested into many different peptides in quantities below the detection limit.

### S4. Circular dichroism experiment

TconTS1 and H-TconTS1 were measured in circular dichroism experiments in a wavelength region from 190 – 250 nm. The secondary structural composition of proteins can be predicted from these spectra by the webserver BeStSeL (60, 61), as final circular dichroism spectra consist of additive signals from secondary structure elements like α-helices or ß-sheets. Table S4 summarizes the extracted secondary structure composition from circular dichroism spectra in Fig. 4A, revealing no significant differences between TconTS1 or H-TconTS1. These results indicate that EndoH_f_ treatment and the removal of *N*-glycans from the protein surface have not influenced the secondary structural composition of H-TconTS1.

**Table S4:** Secondary structure elements of TconTS1 and H-TconTS1, calculated using BeStSeL from the respective circular dichroism spectra (Fig. 4A).

|  | TconTS1 | H-TconTS1 |
| --- | --- | --- |
| $\alpha$ - helix | 13.5 ± 0.2 | 13.2 ± 0.5 |
| β-sheet<br>(antiparallel) | 37.1 ± 2.1 | 36.4 ± 2.5 |
| β-sheet<br>(parallel) | 0 | 0 |
| turn | 11.3 ± 0.1 | 11.2 ± 0.0 |
| others | 38.2 ± 1.9 | 39.2 ± 2.0 |

### S5. Molecular dynamics simulations

Due to missing experimental structures of TS from *T. congolense*, the atomistic structure of TconTS1 was derived using the I-TASSER web server for protein structure and function predictions (63, 64). The engineered SNAP-*Strep* was included for consistency and better comparison with experimental data. The threading algorithm mainly employed TranSA (PDB entry: 2agsA, 2A75) as well as TcruTS (PDB entry: 1ms9) as templates. Validation of the homology modelled TconTS1 was performed by an amino acid sequence alignment of recombinant TconTS1 with TranSA (PDB entry 2ags) and TcruTS (PDB entry: 1ms9) revealing that 10 out of 14 amino acids predicted to be important for enzymatic activities are conserved among all compared models (Fig S6A). Furthermore, 2 of the remaining 4 sites are conserved between TconTS1 and TcruTS (Y211, P379) and only the left over 2 not conserved in TconTS1 (A325, Y408). It needs to be noted that especially Y408, part of the lactose holder pair in the binding site, is a tryptophan in TranSA and TcruTS and therefore both amino acids resemble due to their hydrophobic character. Coloring of the atomistic structure of TconTS1 by the amino acid sequence alignment from Figure S6A is given the impression of most conserved residues to be located in ß-sheet or α-helix regions (Fig S6B). Amino acids of loop regions seem to be less conserved, probably also being less important for the overall structural folding and function of the enzyme. Structural alignment of TconTS1 with TranSA and TcruTS reveal a high similarity of all models with respect to the secondary and tertiary structure (Fig S6C). Only the N-terminal part of TconTS1 is in general longer compared to TranSA and TcruTS and therefore cannot be aligned. Independent prediction of the secondary structure by I-Tasser as well as the inherent thermal mobility of each residue of TconTS1 are akin to that of TranSA and TcruTS (Fig S6D). This is because ß-sheets are dominant in the catalytic and lectin domain, whereas an α-helix is connecting both domains.

Despite an amino acids sequence identity of only 37/38 % between recombinant TconTS1 (with SNAP-*Strep* Tag) and TranSA/TcruTS, the I-Tasser homology model validation suggests a similar secondary structure of TconTS1 compared to other TS from different species and therefore predicts a likewise tertiary model. Conservation of almost all catalytically involved amino acids in TconTS1 further support the idea of structural similarity to the other TS. The I-Tasser confidence score of the recombinant TconTS1 model was given with -2.99 (range -5 to 2), where a higher value signifies a higher confidence. It is, however, necessary to note that the artificial SNAP-*Strep* domain is included in this assessment, and an analysis of only the native TconTS1 results in a confidence score of -0.63, stating a much higher reliability. A confidence score of above -1.5 means that more than 90% of the predictions are correct and therefore our TconTS1 models is considered to be predicted with an overall correct fold (63). Final objections of model accuracy can however only be addressed with structure determination techniques like X-ray crystallography in the future.

**Figure S6:**
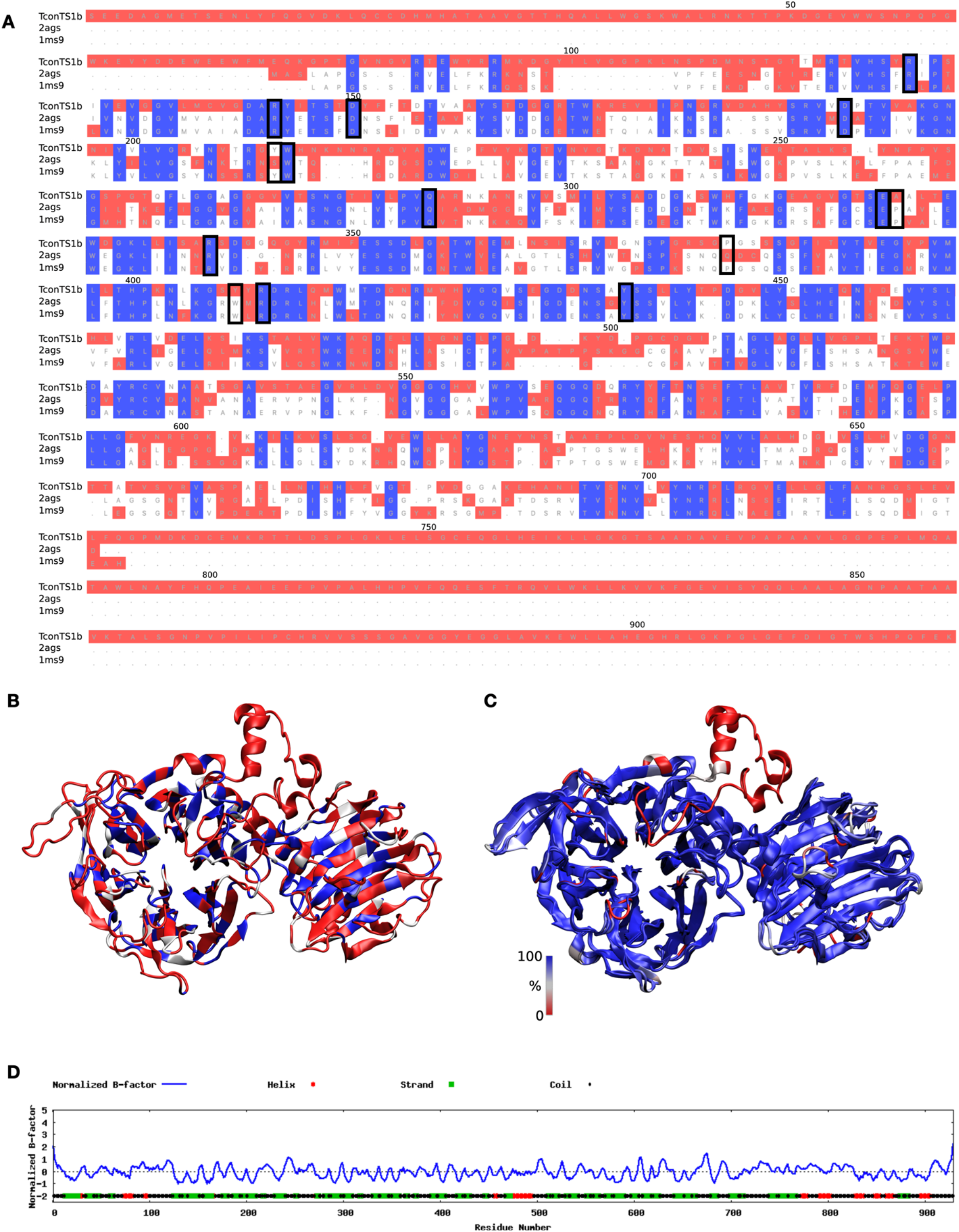
A) Amino acid sequence alignment of recombinant TconTS1, TranSA (PDB entry: 2ags) and TcruTS (PDB entry: 1ms9) by ClustalW using the bioinformatics analysis tool MultiSeq (87) implemented in VMD. Fully conserved amino acids are depicted in blue, partially conserved in white and not conserved in red. Residues of the catalytic domain, which are considered to be important for enzymatic activity are surrounded by a black box (35, 47). Residue numbering is in correspondence with the native TconTS1 sequence. B) 3D structure of TconTS1 in a cartoon style, where coloring of each amino acid is in correspondence with A) The C-terminal SNAP-Strep-Tag is not shown for simplicity. C) Structural alignment of TconTS1, TranSA (PDB entry: 2ags) and TcruTS (PDB entry: 1ms9) by VMD represented in cartoon style. Coloring is in accordance with the Q factor of each residue, where Q is a metric for structural homology implemented in VMD. Blue is referring to 100 % structural identity and a color shift over white to red, a less well alignment. D) Plot of the normalized B-factor, representing the inherent thermal mobility of each residue with indication of predicted secondary structural elements, generated by I-Tasser (63, 64).

Homology modelled TconTS1 was further modified by CHARMM-GUI Glycan Modeler (www.charmm-gui.org) to construct a glycosylated variant termed TconTS1 representing the highest degree of *N*-glycosylation that has been detected by MALDI-TOF MS experiments namely for sites N45, N113, N206, N240, N281 and N693 (Fig S7A). Man_5_GlcNAc_2_ was chosen as a model glycan tree for all sites, as it is the most abundant found form of *N*-glycans in CHO Lec1 cells (41). The EndoH_f_ treated variant was constructed by the attachment of single GlcNAc residues at sites N45, N113, N206, N240, N281 and N693 (Fig S7B), which were previously found to be glycosylated. No remaining high-mannose type *N*-glycan trees are attached to the modelled enzyme, although incomplete deglycosylation has been observed in experiments for certain sites. For consistency, the enzyme is still termed H-TconTS1. It was the aim to represent the highest degree of deglycosylation, by attaching residual GlcNAc at the asparagine residues in N-X-S/T motives.

**Figure S7:**
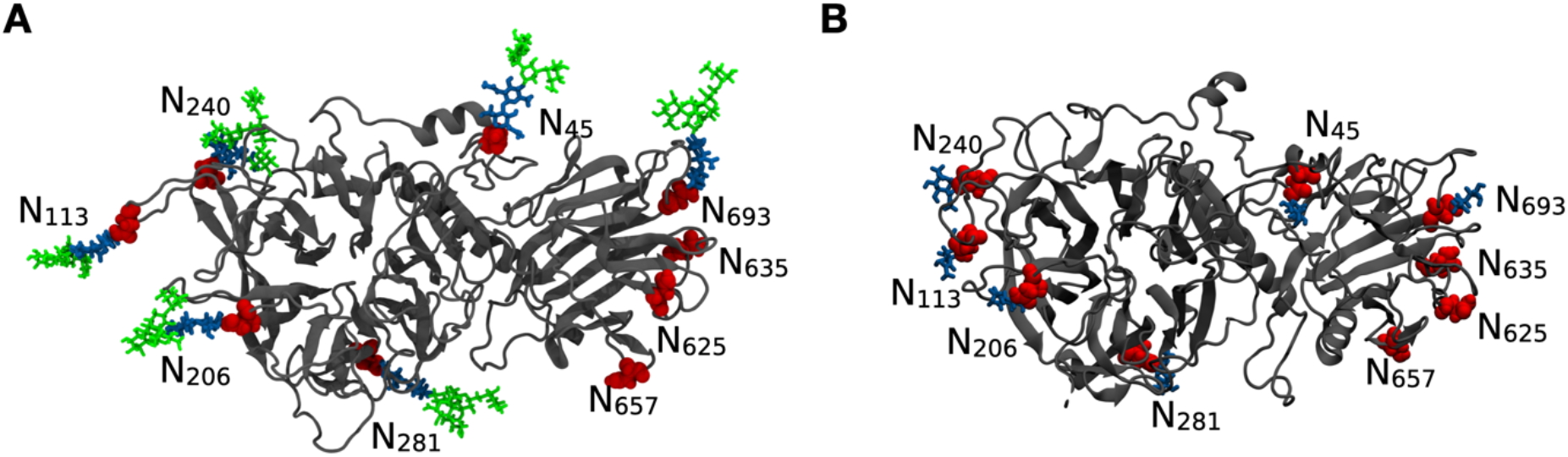
Atomistic structure of TconTS1 A) and H-TconTS1 B) with attached glycan residues to respective *N*-glycosylation sites. Representation and coloring like Figure 5.

Qualitative comparison of secondary structural differences between TconTS1 and EndoH_f_ treated H-TconTS1 were not only addressed experimentally by circular dichroism experiments, but also further verified by MD simulations. Therefore, the assignment of secondary structure elements was tracked over time for each residue (Fig. S8). In total, it can be said that the overall secondary structure of TconTS1 and H-TconTS1 is comparable over the simulated time, as α-helices in the N-terminal region (position 67 – 76), between the catalytic and lectin domain (position 468 – 489) as well as in the C-terminal SNAP-*Strep* region (position from 707 onwards, Fig. S1A) are consistent. The same applies to the shorter but more frequently observed ß-sheet regions. These simulated results support the findings of circular dichroism experiments, in which no changes in secondary structure elements could be observed, too.

Additionally, it can be seen that *N*-glycosylation sites are mostly situated in coil and turn secondary structural motifs, positioned next to ß-sheet regions. The same applies to residue D150, positioned in-between two ß-sheet regions, with at least 5 amino acids assigned to coil and turn elements directly next to each other. These observations hint at a certain flexibility of *N*-glycosylation sites and amino acid D150, not involved in extensive secondary structure motives of α-helices or ß-sheets.

**Figure S8:**
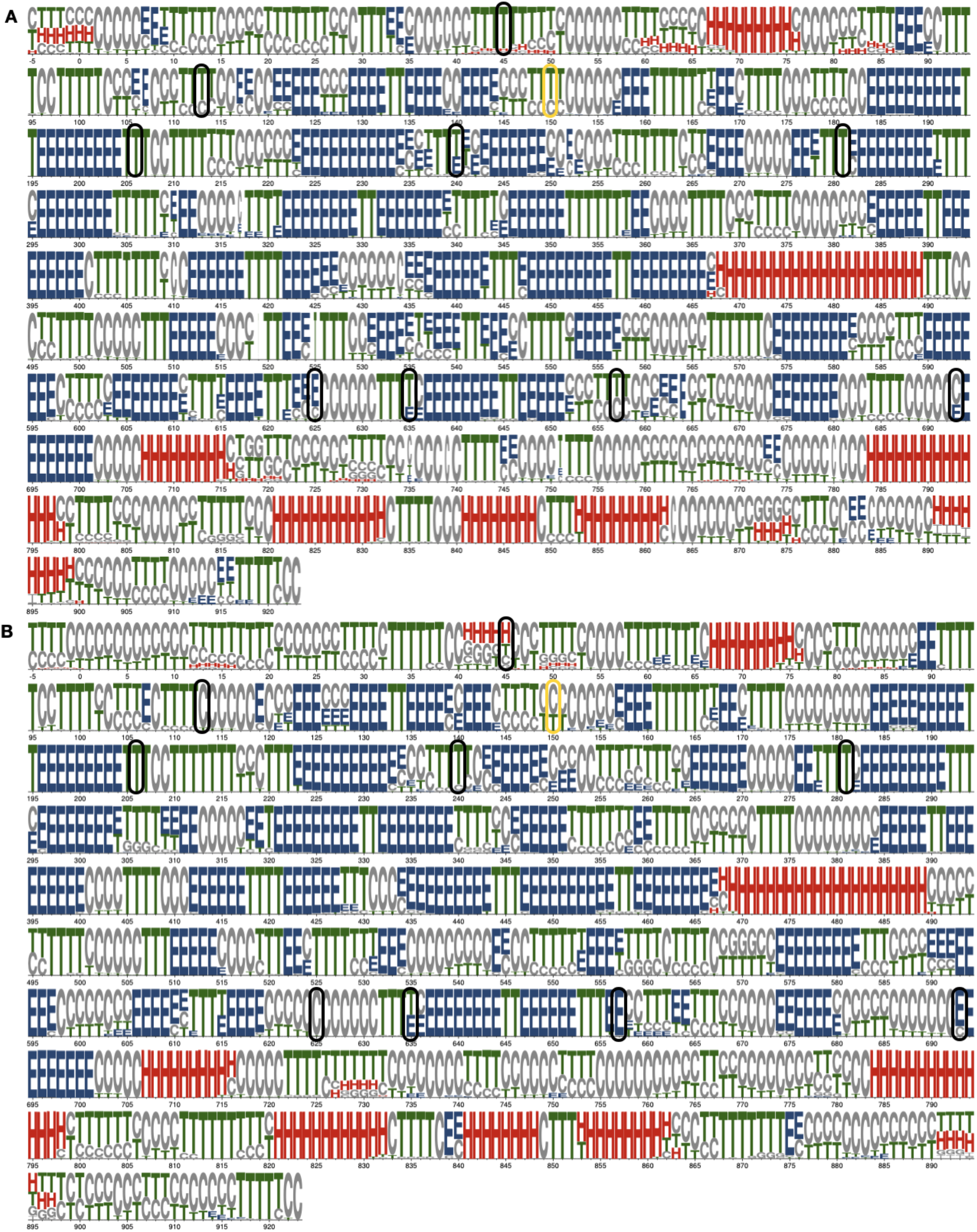
A) Secondary structure of TconTS1 averaged over 500 ns of classical MD. B) Secondary structure of H-TconTS1 averaged over 500 ns of classical MD. Size of letters is corresponding to the probability of secondary structure element occurrence. H = α-helix (red), E = ß-sheet (blue), T = turn (green), C = coil, B = Isolated bridge, I = Pi-helix (grey). Positions of *N*-glycosylation sites are marked with black circles. Position of D150 is marked in yellow. Secondary structure was calculated using the STRIDE module of VMD (88). The figure has been created using the weblogo 3 website (89).

